# DNA barcoding reveals ongoing immunoediting of clonal cancer populations during metastatic progression and in response to immunotherapy

**DOI:** 10.1101/2021.01.11.426174

**Authors:** Louise A. Baldwin, Nenad Bartonicek, Jessica Yang, Sunny Z. Wu, Niantao Deng, Daniel L. Roden, Chia-Ling Chan, Ghamdan Al-Eryani, Damien J. Zanker, Belinda S. Parker, Alexander Swarbrick, Simon Junankar

## Abstract

Cancers evade the immune system in order to grow or metastasise through the process of cancer immunoediting. While immune checkpoint inhibitors have been effective for reactivating tumour immunity in some types of cancer, many other solid cancers, including breast cancer, remain largely non-responsive. Understanding the way non-responsive cancers evolve to evade immunity, what resistance pathways are activated and whether this occurs at the clonal level will improve immunotherapeutic design. We tracked cancer cell clones during the immunoediting process and determined clonal transcriptional profiles that allow immune evasion in murine mammary tumour growth in response to immunotherapy with anti-PD1 and anti-CTLA4. Clonal diversity was significantly restricted by immunotherapy treatment in both primary tumours and metastases. These findings demonstrate that immunoediting selects for pre-existing breast cancer cell populations and that immunoediting is not static, it is ongoing during metastasis and immunotherapy treatment. Isolation of immunotherapy resistant clones revealed unique and overlapping transcriptional signatures. The overlapping gene signature was associated with poor survival of basal-like breast cancer patients in two cohorts. At least one of these overlapping genes has an existing small molecule that can potentially be used to improve immunotherapy response.

## Introduction

All cancers must find ways to evade the immune system so that they can continue to grow (*1*). Previous studies have established that this occurs through a process called immunoediting (*2*). During immunoediting, more immunogenic cancer cells are selectively eliminated by the immune system thus leaving behind less immunogenic cancer cells that are then free to expand. Immunoediting can occur through multiple mechanisms, which include the elimination of cells with strong immunogenic mutations, leading to the loss of neo-antigens (*3*), or the selection of cells with elevated expression of various immunosuppressive programs (*4*).

Immunotherapies look to overcome some of the immune evasion pathways established by cancer cells. The prominent clinically approved immunotherapies for solid tumours target T cell checkpoint molecules (eg. anti-CTLA4 and anti-PD1) to overcome T cell exhaustion (*5, 6*). In select cancer types, such as melanoma, immune checkpoint inhibitors have dramatic effects in a large proportion of patients (*7*). Unfortunately for metastatic breast cancer, few patients, even those having the most sensitive basal-like breast cancer, had durable responses in clinical trials (*8*). This indicates that in metastatic breast cancer, resistance is either pre-existing or can rapidly develop to anti-PD1/PDL1 therapy and suggests that alternate immune drug targets are needed for breast cancer.

Although the immune system is known to play a role in breast cancer outcome (*9*) and immunoediting can occur in a transgenic mouse model of breast cancer (*10*), very little is known about immune evasion by breast cancer cells. The majority of studies examining immune evasion by cancers were performed using the highly mutated, methylcholanthrene-driven sarcoma model, the response of which cannot be tracked at the clonal level (*11*), or colon cancer (*12*). Of interest, a recent study suggests that immunoediting by T cells can occur at the clonal level by demonstrating the selection of clones that contain less-immunogenic fluorophores (*13*). This leaves an important gap in our collective knowledge as to the mechanisms employed in less immunogenic tumours such as breast.

Both natural killer (NK) cells and T cells have been demonstrated to play a role in immunoediting (*14, 15*)). However, the majority of recent research has focussed on pathways relevant to T cell recognition (*16–20*). Downregulation of MHC is one mechanism by which cancer clones become impervious to T cells (*21*), but this inherently makes them targets of NK activity. In breast cancer dysfunction of NK cells is noted and this is regulated by microenvironmental factors (*22*). Data on resistance pathways that allow for immune evasion from both T cells and NK cells are currently more limited.

Intratumoural heterogeneity (ITH) has been identified as a major contributor to treatment response. Prior work in non-small cell lung cancer (NSCLC) demonstrated that neoantigen heterogeneity, and tumour mutation burden more broadly, is strongly associated with T cell anticancer immune responses (*23, 24*). Chemotherapy(*25*) or loss of HLA (*26*) have also been shown to increase ITH, which in turn was associated with treatment relapse in NSCLC. Similar results have been described in cell line models of melanoma (*27*), with Williams and colleagues providing recent evidence of ITH enabling clonal cooperatively and immune escape in melanoma-derived cell lines (*28*). While these previous studies have been crucial to our understanding of ITH and anti-tumour immune responses and have provided undeniable evidence associating increased ITH with therapy resistance, studies of this nature have not yet been carried out in breast cancer. Furthermore, direct evidence of ITH enabling immune escape and immunotherapy resistance in vivo has not been reported.

DNA barcoding allows tracking of individual clones and clonal expansion while avoiding introducing potentially immunogenic proteins (*29*). We used DNA barcoding to analyse immunoediting *in vivo* at primary tumours and metastases and to study whether resistance to immune checkpoint inhibition develops from pre-existing or *de novo* generated cell populations.

## Results

### Immunoediting of breast cancer cells in the primary tumour

To understand the role of the immune system and immunotherapy in shaping the clonal dynamics of cancer cells within primary tumours, we used the DNA barcoding approach (Fig 1). We introduced the ClonTracer DNA barcode library that contains approximately 7 million unique barcodes (*30*) into the immunotherapy-sensitive mouse mammary carcinoma EMT6 cells (*31*) (Sup Fig 1), resulting in ∼41 000 unique barcodes identified by DNA sequencing (Sup Fig 2A) . We then inoculated 250 000 cells (∼6 fold over representation of each barcode) into the mammary fat pad of syngeneic immune-competent wild-type (WT) Balb/c mice or severely immunocompromised NOD SCID Gamma (NSG) mice that lack T cells, B cells and functional NK cells, and compared the number of clones that were able to engraft and grow (Fig 2A, Sup Fig 2).

**Figure 1.**
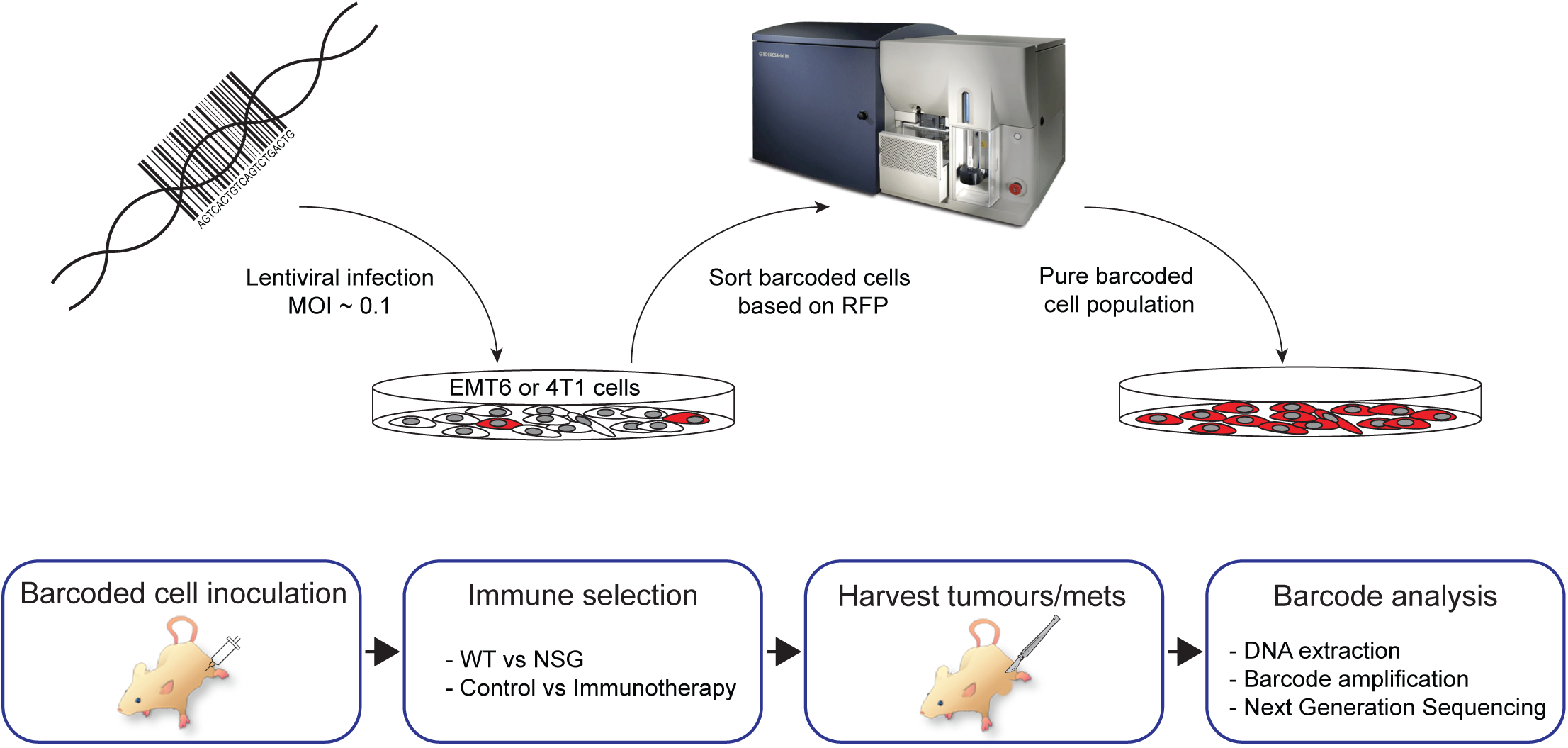
Experimental workflow schematic. Barcode library is introduced into mammary carcinoma cell lines *in vitro* at a low multiplicity of infection (MOI). Cells are sorted based on RFP expression to select for those having incorporated a barcode. Barcoded cells are then transplanted into the mammary fat pad of mice. Following immunoselection with either the endogenous immune system or immunotherapy then barcode abundance and diversity within the primary tumours and lungs with metastases are analysed.

**Figure 2.**
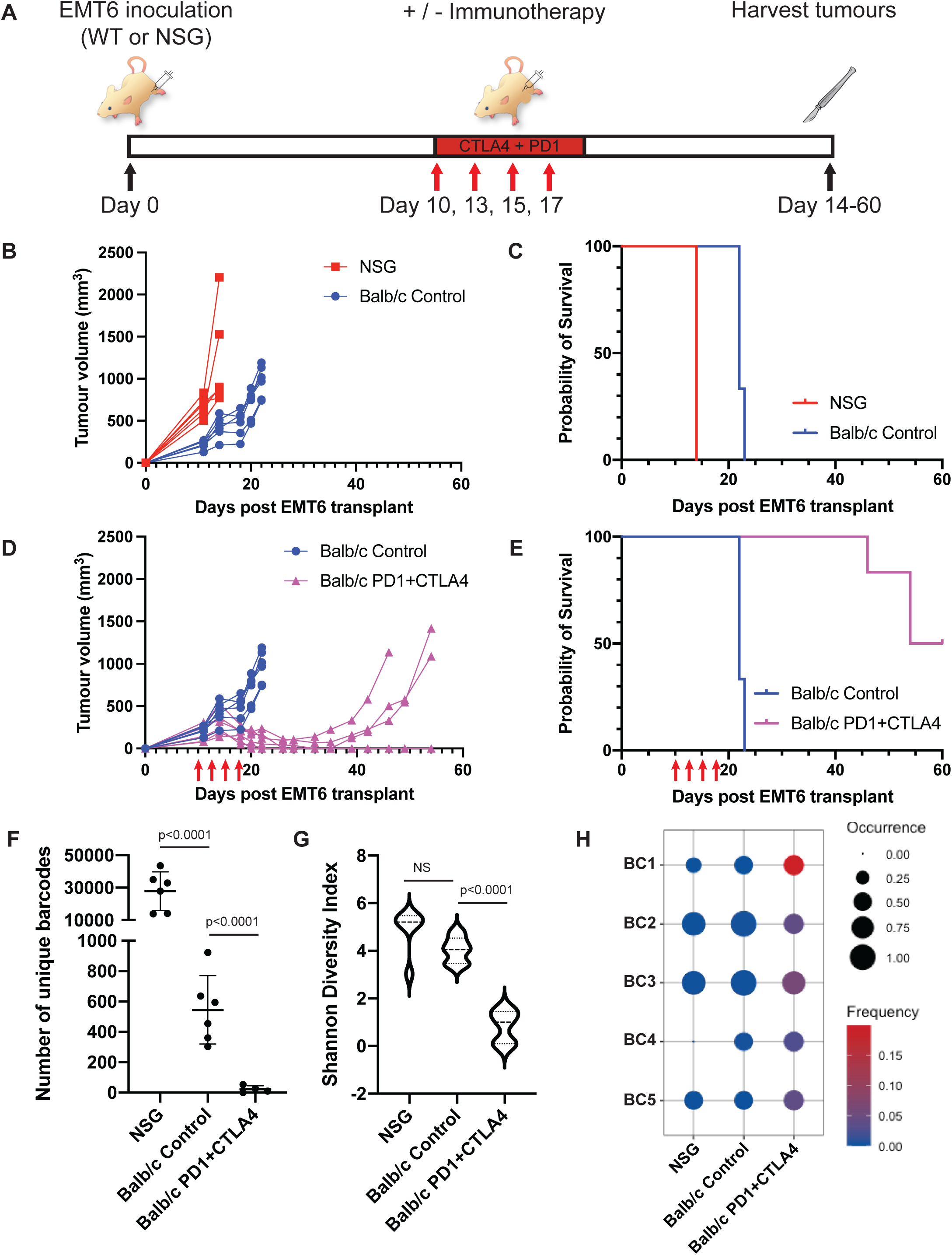
Immune selection and clonal immunoediting of EMT6 primary tumours. A. Outline of experimental design. B. EMT6 primary tumour growth in wild-type Balb/c mice and in NSG mice plotted as tumour volume. Average volume +/- SEM; n=5-6 mice per group. C. Kaplan-Meier survival analysis comparing Balb/c and NSG mice bearing EMT6 tumour (Mantel-Cox p=0.009). D. EMT6 primary tumour growth plotted as tumour volume in wild-type Balb/c mice with or without immunotherapy (anti-PD1+anti-CTLA4) on days 10, 12, 14, and 17 as indicated with red arrows. Average volume +/- SEM; n=5-6 mice. E. Kaplan-Meier survival analysis of Balb/c mice bearing EMT6 tumour treated with immunotherapy or isotype control (Mantel-Cox p=0.0006). F. Number of unique barcodes identified in EMT6 primary tumours grown in NSG mice and in Balb/c mice treated with isotype control antibodies or anti-PD1 + anti-CTLA4. GLM with Tukeys HSD for multiple comparisons. G. Shannon diversity index analysis of EMT6 primary tumours grown in NSG mice and in Balb/c mice treated with isotype control antibodies or anti-PD1 + anti-CTLA4. One way ANOVA with Tukeys HSD. H. Dot plot of a subset of barcodes with enrichment following immunotherapy treatment.

Tumour growth was faster in NSG mice than in WT mice, with tumours reaching ethical endpoint in NSG mice on day 14 post-transplant and in WT mice by day 23 (Fig 2B), leading to NSG mice having significantly shorter median overall survival (14 days) than WT mice (22 days, Mantel-Cox p=0.009). These results suggest that the immune system plays an important role in controlling the growth of the EMT6 primary tumours (Fig 2C).

To examine the influence of immunotherapy on tumour growth and clonal dynamics, we compared WT mice treated with immunotherapy (anti-PD1 + anti-CTLA4) or control antibodies starting from day 10 when tumours were approximately 200mm^3^ (Fig 2A). Tumours in all mice treated with control antibodies reached ethical endpoint by day 23 (Fig 2D). In contrast, tumours in all mice treated with immunotherapy regressed following treatment, with 50% relapsing and reaching ethical endpoint between days 46 and 54 (Fig 2D). The remaining mice treated with immunotherapy remained tumour free until the experiment was terminated on day 60. During harvest a small residual lesion was observed in two of these mice, no metastatic lesions were observed in any of the mice irrespective of treatment. Kaplan-Meier analysis demonstrated that immunotherapy significantly increased the median survival from 22 days to 57 days (Mantel-Cox p=0.0006) (Fig 2E).

To determine if immune control of tumour growth was driven at a clonal level, we examined the number and distribution of barcodes present in primary tumours collected at ethical endpoint in the experiment described above. We found that at ethical endpoint, tumours grown in NSG mice had over 50 times the number of unique barcodes as tumours grown in control WT mice (p<0.0001, generalised linear model (GLM) with Tukey’s correction), which in turn had more than 20 times the number of unique barcodes found in WT mice treated with immunotherapy (p<0.0001, GLM with Tukey’s correction) (Fig 2F, Sup Fig 2A). We applied Shannon diversity analysis to understand how the immune system influenced the diversity of barcodes in these samples. Shannon diversity index determines how evenly distributed the barcodes are within a population and is only moderately influenced by barcode number. Analysis of barcode diversity revealed a trend to a lower barcode diversity in tumours from control (WT Balb/c) mice than that in tumours from NSG mice whereas the barcode diversity was significantly lower in immunotherapy treated than in control treated Balb/c mice (Fig 2G, p<0.001 one way ANOVA with Tukey’s correction). These data suggests that a subset of EMT6 cells are more resistant to the endogenous immune system but this selection does not skew the evenness of the barcode distribution significantly, which suggests that all clones resistant to the immune system have similar levels of resistance. In contrast, immunotherapy applies a more stringent bottleneck that only a limited number of clones can overcome, and with a high degree of variability in doing so. Further analysis identified EMT6 clones that were reproducibly enriched across multiple mice following immunotherapy treatment, indicating that they had a pre-existing resistance phenotype that was being positively selected for (Fig 2H).

### Immunoediting of breast cancer cells during metastasis

To determine whether immunoediting continued during metastatic dissemination and whether specific metastatic clones were enriched or depleted, we turned to the highly metastatic 4T1 mammary carcinoma model, as the EMT6 cell line is poorly metastatic (*32*). We introduced the barcode library into 4T1 cells, leading to a cell pool with ∼5000 unique barcodes (Sup Fig 3, Sup Fig 4), then inoculated 50,000 of these cells (a ∼ 10-fold over-representation of each barcode) into the mammary fat pad of WT and NSGmice (Sup Fig 4). We resected primary tumours 15 days following inoculation to allow metastases to develop (Fig 3A). All mice developed lethal lung metastases, with NSG mice succumbing to metastatic disease earlier than WT mice (median survival of 25.5 days verses 35 days, p = 0.0002, Mantel-Cox Log-rank test) (Fig 3B). Primary tumour sizes at the time of resection were similar between the groups (Sup Fig 5). Adjuvant immunotherapy with anti-PD1 + anti-CTLA4 led to a modest but significant increase in survival (37.5 days) versus control-treated mice (33 days; p=0.0121, Mantel-Cox Log-rank test) (Fig 3C).

**Figure 3.**
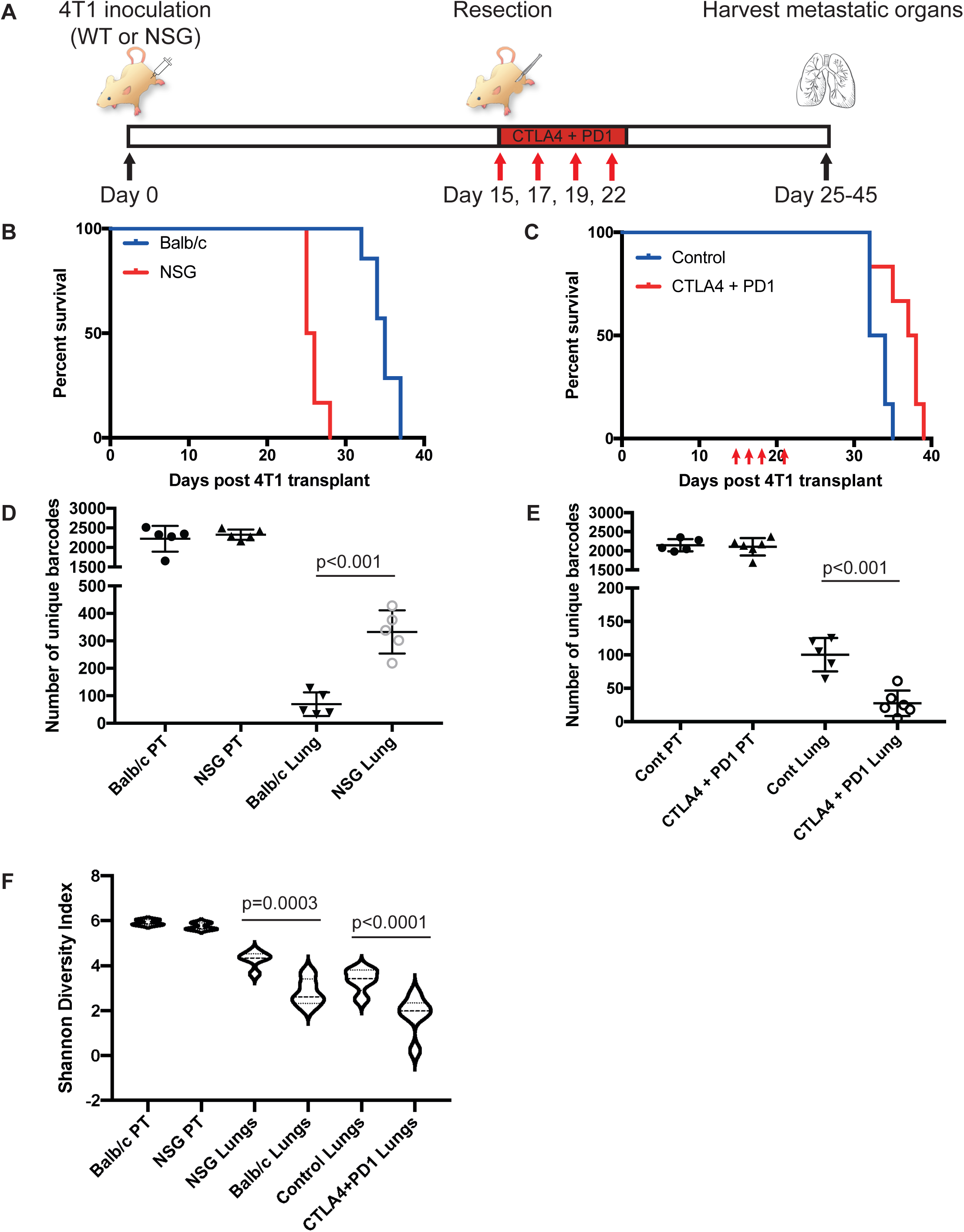
Immune selection and clonal immunoediting in the metastatic setting utilising the 4T1 model. A. Outline of experimental design. B. Kaplan-Meier survival analysis comparing Balb/c mice and NSG mice bearing 4T1 tumour; n=5 mice/group (p = 0.0002, Mantel-Cox Log-rank test). C. Kaplan-Meier survival analysis of Balb/c mice bearing 4T1 tumour treated with immunotherapy or isotype control on days 15, 17, 19, and 21 as indicated with red arrows; n=5-6 mice/group (p = 0.0141, Mantel-Cox Log-rank test). D. Number of unique barcodes identified in 4T1 primary tumours (PT) and lung metastases grown in NSG mice or Balb/c mice. GLM with Tukey’s HSD for multiple comparisons. E. Number of unique barcodes identified in 4T1 primary tumours and lung metastases grown in Balb/c mice treated with isotype control antibodies or anti-PD1 + anti-CTLA4. GLM with Tukey’s HSD for multiple comparisons. F. Shannon diversity index analysis in primary tumours from NSG mice or Balb/c mice and in lung metastases from NSG mice, Balb/c mice, Balb/c mice treated with isotype control, and Balb/c mice treated with anti-PD1 + anti-CTLA4. Unpaired t-test. G. Analysis of number of unique barcodes in lung metastases from Balb/c mice treated with control antibodies, anti-CD8a to deplete CD8 T cells, or anti-asialo GM1 to deplete NK cells. One way ANOVA with Tukeys HSD for multiple comparisons, 5 mice/group.

We then examined whether the endogenous immune system shaped metastatic clonal dynamics. While primary tumours contained similar numbers of clones and barcode diversity in NSG and WT hosts (Fig. 3D and Sup Fig 5B, C), the lung metastases of NSG mice contained ∼3 times as many barcode clones as those of WT controls (Fig 3D). We next determined if the increase in survival following immunotherapy was associated with alterations in clonal dynamics. As the treatment was only given after excision of the primary tumour, the immunotherapy would only affect the outgrowth of cancer cells that had already metastasised to the lung. Despite only a modest increase in survival following combination immunotherapy (Fig 3C), we observed a 70% reduction in the number of clones in metastases (Fig 3E).

The higher barcode number in the lung metastases of NSG mice than in the lung of WT mice was associated with a higher diversity of barcode as measured using the Shannon diversity index (Fig 3F). This shows that the endogenous immune system restricts the number and skews the diversity of metastatic clones that can reach and outgrow in the lungs. In addition to the reduction in barcode number following immunotherapy treatment, we also saw a significant reduction in barcode diversity (Fig 3F). This suggests that immunotherapy is leading to the immunoediting of specific clonal cell populations over others.

To further understand the key immune cell types that control clonal outgrowth in the lung, we depleted CD4 T cells using anti-CD4, CD8 T cells using anti-CD8 or NK cells using anti-asialo-GM1 in wildtype mice starting one day prior to tumour resection (Sup Figs 6B, 6C, 6D). None of these treatments led to a significant change in overall survival (Sup Fig 6A). Initial experiments indicated a small change in the number of clones detected within the lung between treatment groups (Sup Fig 6E). However, this was not recapitulated in repeat experiments (Sup Fig 6F). This may suggest that none of these cell types alone are sufficient to restrict clonal diversity after seeding of pulmonary metastases.

We were surprised with the lack of immunoediting in the primary tumours of the 4T1 model so we repeated these experiments with a second pool of barcoded 4T1 cells containing a larger barcode library (300 000 barcodes). Following the injection of 50 000 barcoded cells, we recovered approximately 10 000 - 12 000 barcode sequences from each primary tumour grown in WT mice and in NSG mice (Sup Fig 8A). This suggests that roughly a fifth of the injected cells are able to engraft and grow in the mammary gland. As clone diversity was again similar between NSG mice and WT mice, this confirms that the immune system does not play a major role in restricting growth of 4T1 cells in the primary tumour setting. In contrast, when we examined the number of clones that had spread to the lung we again found approximately 3 times as many in NSG mice as in wild-type mice (Sup Fig 8A). In addition, we similarly saw a ∼3-fold reduction in barcode diversity in response to immunotherapy (Sup Fig 8B). These results confirm that the 4T1 cell line is already highly immunoedited for growing at the primary site yet is subjected to a second round of immunoediting during metastasis.

### Patterns of enrichment and depletion of specific clones

To better understand how specific clonal cell populations responded to the immune system and immunotherapy, we combined the barcode frequencies from the two datasets utilising the 5000 barcodes library (WT vs NSG and Control vs Immunotherapy). We performed unsupervised hierarchical clustering of these samples and selected barcodes that were observed at greater than 5% frequency in any one sample. We found that the primary tumours from the two experiments cluster together irrespective of the immune status of the mouse (Balb/c or NSG), further suggesting that 4T1 primary tumours, in contrast to EMT6 tumours, do not undergo further immunoediting (Fig 4A). In contrast, lung tumours formed in the NSG hosts did not cluster with lung tumours formed in Balb/c mice, with the immunotherapy treated samples mostly clustering alone or with metastases formed in WT mice. A number of specific barcodes were enriched in lung metastases of all NSG mice, indicating that these clones were highly metastatic (Fig 4A), this is similar to the findings of Wagenblast and colleagues who examined 4T1 clonal diversity in metastases in NSG (but not wild type mice) and found tissue-specific enrichment of unique barcode clones(*33*).

**Figure 4.**
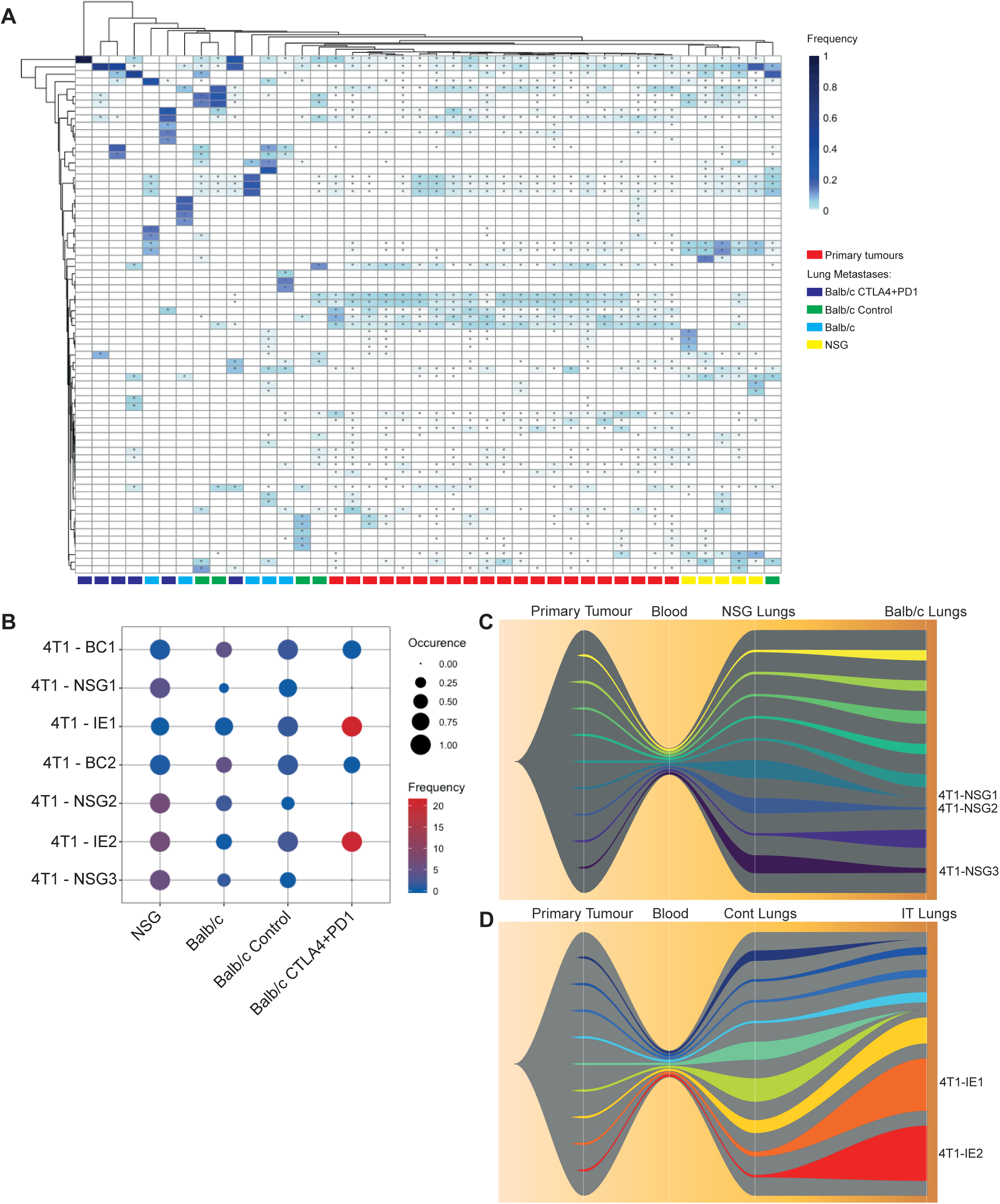
Analysis of specific barcodes enriched or depleted by the immune system and immunotherapy in the 4T1 model. A. Unsupervised hierarchical clustering heatmap of barcodes with an abundance of above 5% in at least one 4T1 primary tumour or lung metastasis sample; * indicate barcodes detected at a frequency above 0.1% in a particular sample. B. Dot plot of a subset of specific barcodes. C. Fishplot of the nine most abundant barcodes detected in lungs of WT and NSG mice, each of these nine barcodes is given a unique colour with the remaining barcodes being combined and represented in grey. A bottleneck has been introduced between primary tumour and lung metastases to depict transition through the blood stream. D. Fishplot of the nine most abundant barcodes detected in the lungs of WT mice treated with combination immunotherapy or control antibodies presented as in C, a bottleneck has been introduced between primary tumour and lung metastases to depict transition through the blood stream.

We identified from the heatmap (Fig 4A) a number of barcodes that had striking patterns of enrichment or depletion in response to the immune system and immunotherapy, we replotted these using a dot plot (Fig 4B). These barcodes were enriched or depleted in a reproducible manner across replicate mice, suggesting this is due to inherent features of these clones. There are three barcodes that were enriched in NSG lung metastases (NSG1-3), present at lower abundance in untreated wild-type mice, and were completely eliminated following immunotherapy treatment. This suggests that these clones are highly metastatic in the absence of an immune system but they are immunogenic and are thus subjected to immunoediting in WT mice, particularly following immunotherapy. Another group of metastatic clones present in the lung of NSG and WT mice were further enriched following immunotherapy (IE1-2). These immunotherapy enriched clones were detected in the lungs of all six replicate mice. With the significant reduction in the number of barcodes present following immunotherapy, the odds ratio of this reproducible enrichment happening by chance is 0.0034 (95% confidence interval: 0.0010-0.0079; chi square p value: 3.67x10^-251^). This suggests that these clones have a pre-existing resistance phenotype and are positively selected for following immunotherapy.

To further analyse how specific barcodes were enriched in lung metastases following immunotherapy, we visualised the top nine clonal populations (based on average barcode proportions in the metastatic lungs) and generated fish-plots. These showed that different clones were preferentially enriched in the lung of NSG mice when compared to WT mice (Fig 4C). Furthermore, we observed that a small subset of clones was highly enriched in the lungs of immunotherapy treated mice (Fig 4D).

### Analysis of immunotherapy resistant clones

To understand more about the phenotype of these immunotherapy resistant clones, we established clonal cell populations from two of them (designated IE1 and IE2) and from two independent control clones (NT1 and NT2) that were not enriched following immunotherapy. We generated 3-4 independent clonal cell lines per barcode clone. These clonal cell lines were isolated from the parental barcoded 4T1 cell population *in vitro* with no additional selective manipulation. The barcode within each of these clonal cell lines was confirmed to be correct using Sanger sequencing. All four clonal cell lines had similar growth kinetics *in vitro* indicating no proliferative advantage of the immune evasive clones *in vitro* (Sup Fig 9A). All clonal cell lines were able to form tumours in the primary setting (Sup Fig 9B). IE2 demonstrated considerable metastatic potential with 66% of mice forming extensive lung metastases and regularly metastasised to the lungs (Sup Fig 9C). IE1 also demonstrated some metastatic capacity with 30% of mice forming lung metastases. Neither NT1 nor NT2 successfully metastasised to the lungs (Sup Fig 9C).

### Genomic analysis for barcode integration site and copy number variation (CNV)

To identify the barcode integration sites and determine whether the clones contained large scale genomic alterations, we performed whole genome sequencing (WGS) at around 30x coverage of the clones. The WGS analysis determined the precise genomic location where barcodes integrated (Sup Table 2). The integration site in IE1 was in the intergenic region between *Kpna*2 and *Smurf*2 and the integration site in IE2 was within an intron of *Nrf*1, neither integration site changed the coding sequence of these genes. Copy number analysis determined that no clone had dramatic copy number changes when compared to other clones. Each clone only contained a small number of single copy number gains and losses (IE1 only 6 CNVs and IE2 only 5 CNVs), with clone NT2 showing the greatest number of CNVs at 41 (summarised in Sup Table 3). We found one locus on chromosome 18 with a single copy number gain in both IE1 and IE2 that led to 3 copies of *Nc3r1*, encoding the Glucocorticoid receptor, and *Arhgap26*, encoding a Rho GTPase that associates with focal adhesion kinase. However, this copy number gain on chromosome 18 was also present in the NT2 clone. These results demonstrate that large scale genomic changes unlikely play a major role in determining various phenotypes of different clones but instead suggest that copy number changes may be selected against during immunoediting.

### Transcriptomic analysis of the clonal cell lines

To investigate the mechanism of immune evasion by these clones, we performed RNAseq analysis and compared the two immunotherapy resistant clones to the bulk 4T1 population. Differential gene expression analysis was carried out using EdgeR. The IE1 clone had 1553 differentially expressed genes (Log fold change >2 and FDR p<0.05) with 478 significantly upregulated and 1075 significantly downregulated (Fig 5A). The IE2 clone had 1099 differentially expressed genes with 375 significantly upregulated and 724 significantly downregulated (Fig 5B). The non-target clones had fewer gene expression changes compared to the bulk 4T1 population with NT1 having 621 and NT2 having only 262. We examined the top differentially expressed genes between each of IE1 and IE2 with the parental 4T1 cells, however, we did not find any with an obvious role in immune evasion (Sup Tables 4 & 5). Gene set enrichment analysis revealed that among the top ten gene sets upregulated in IE1 and in IE2, only two (CHEN_HOXA5_TARGETS_9HR_UP, BLUM_RESPONSE_TO_SALIRASIB_UP) were common between them (Sup Tables 6 & 7). *Hoxa5* is a known tumour suppressor gene in breast cancer (*34*), although we see an enrichment of its target genes, the expression of *Hoxa5* itself was significantly reduced in the IE1 clone and trended to be reduced in the IE2 clone. There was no overlap in the top ten downregulated gene-sets between IE1 and IE2. The top downregulated gene-set for IE1 was the REACTOME_UB_SPECIFIC_PROCESSING_PROTEASES gene-set, which contained two genes involved in antigen processing for display by MHC-I (*Psmb8* and *Psmb9*). As down regulation of the MHC-I pathway is a common mechanism of immune evasion, we investigated this in more detail.

**Figure 5:**
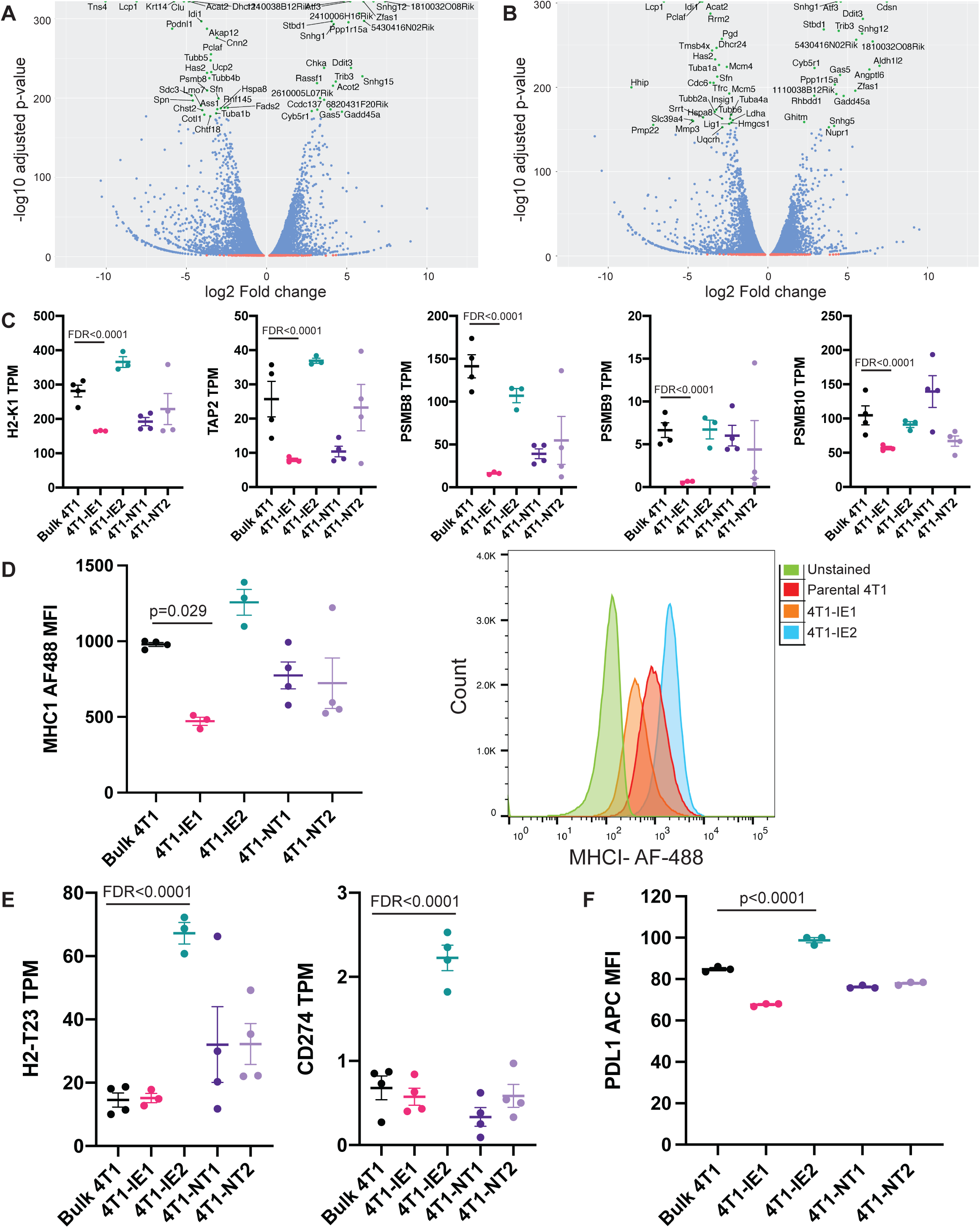
Gene expression analysis of immunotherapy resistant clones. A. Volcano plot showing differentially expressed genes between parental 4T1 cells and the immunotherapy enriched 1 (IE1) clone. B. Volcano plot showing differentially expressed genes between parental 4T1 cells and the immunotherapy enriched 2 (IE2) clone. C. Expression of indicated MHC related genes, measured as transcripts per million (TPM), in the parental 4T1 population and in indicated cell clones; FDR calculated using EdgeR with Benjamini-Hochberg multiple testing correction. D. MHCI protein expression as quantified by flow cytometry in the indicated clones and the parental 4T1 population measured as mean fluorescence intensity (MFI) (left) and representative histogram (right); one way ANOVA with Tukey HSD test for multiple comparisonst. E. Expression of immune related genes up-regulated in clone IE2, measured as transcripts per million (TPM), in the parental 4T1 population and in indicated cell clones. FDR calculated by EdgeR with Benhamini-Hochberg multiple testing correction F. PD-L1 protein expression as determined by flow cytometry in indicated clones and the parental 4T1 population measured as mean fluorescence intensity (MFI), representative plot of three independent experiments; one way ANOVA with Tukey HSD for multiple comparisons.

Through this analysis we found that the IE1 clone had significantly reduced expression of many genes related to antigen presentation, including MHC-I (*H2-k1*), *Tap2*, *Psmb8*, *Psmb9* and *Psmb10* (Fig 5C). *H2-k1* encodes the main MHC molecule that is expressed by the Balb/c mouse from which the 4T1 carcinoma cell line was derived from. We validated the reduction in MHC-I expression levels seen in the RNAseq data at the protein level using flow cytometry (Fig 5D). This analysis showed that the IE1 clone had significantly reduced cell surface MHC-I protein compared to the bulk 4T1 population. We thus examined the WGS data and found that the loss of MHC-I expression in IE1 was not due to genomic loss at the MHC locus on chromosome 17 (Sup Fig 10A). In contrast, the IE2 clone had elevated levels of a number of these MHC related genes (Fig 5C) in addition to *H2-t23* that encodes a non-classical MHC molecule (Fig 5E) known to negatively regulate NK cells through their inhibitory receptor NKG2A (*35*). Interestingly, IE2 cells also demonstrated a significantly increased expression of *Cd274* that encodes the T cell inhibitory molecule PD-L1 (Fig 5E). This again was validated at the protein level using flow cytometry (Fig 5F). These results demonstrate that these two immunotherapy resistant clones are phenotypically unique. To determine if these two mechanisms are simultaneously maintained *in vivo*, we carried out flow cytometry for MHC-I and PD-L1 on advanced 4T1 lung metastases from immunotherapy treated BALB/c mice. By isolating RFP+ cancer epithelial cells from lung tissue, we observed MHC-I-high and MHC-I-low neoplastic populations, both with varying expression of PD-L1 (Sup Fig 11). This provides convincing evidence of these two mechanisms of immune evasion being maintained simultaneously *in vivo*.

We next examined whether copy number changes or barcode integration sites identified above impacted gene expression. The copy number changes of *Nc3r1* and *Arhgap26* were associated with significantly increase of their expression in IE1 and IE2 cells but not in NT2 cells (Sup Fig 10B). Elevated *NC3R1* expression has been associated with poor prognosis and metastasis in triple-negative breast cancer, although whether it plays a role in immune evasion is not known (*29, 30*). As stated above, the barcode integrated in the intergenic region between *Kpna2* and *Smurf2* in IE1 and within an intron of *Nrf1* in IE2. Among these three genes, only the expression of *Smurf2* was significantly altered with a modest log fold increase of 0.59 in IE1.

### Demethylating drugs do not fully restore MHC expression

Demethylating agents such as 5-aza-2’-deoxycytidine (5-aza) are known to upregulate MHC-I expression in cancer cells (*36*), thus we treated our clonal cell lines with 5-aza for 72 hours to determine whether DNA methylation was a mechanisms suppressing MHC expression in the IE1 clone. Using flow cytometry, we observed that MHC-I expression was elevated in a dose dependent manner following 5-aza treatment in all clones. However, MHC-I expression in the IE1 clone was consistently lower than the parental 4T1 cell line at all doses of 5-aza (Sup Fig 12A). This indicates that gene hyper-methylation is not the mechanism of MHC-I suppression in the IE1 clone.

Interferon (IFN)-gamma(g) stimulation is another mechanism by which MHC expression can be increased on cancer cells. The IE1 clone responded to IFN-gamma treatment by upregulating MHC-I expression but again it remained suppressed compared to the parental 4T1 cells (Sup Fig 12B). This suggests these cells broadly retain the transcriptional regulatory machinery that is required to upregulate MHC-I in response to IFN-gamma stimulation. These results indicate that MHC-I downregulation is likely regulated by epigenetic factors other than DNA methylation and that the majority of MHC-I expression in this clone can be restored by IFN-g treatment.

### The 4T1-IE2 clone can directly suppress anti-cancer CD8 T-cell responses

To further characterise the immunotherapy resistant clones (IE1 and IE2) in comparison to the control (NT1 and NT2) clones we performed an *in vitro* CD8 T cell activation assay (*37, 38*). This assay utilises a pool of *de novo* generated 4T1 specific CD8+ T cells, with intracellular IFN-γ production utilised as a functional measure of T cell activation. We then determined how T cell activation changed following co-culture with the clonal 4T1 populations. In the absence of 4T1 cells 1% of CD8 T cells had detectable intracellular IFN-γ production (Fig 6A) while in the presence of the parental 4T1 cells 23% of the T cells had detectable intracellular IFN-g production. When the activation of T cells by cloal cell lines was examined, only the IE2 clone significantly reduced CD8 T cell activation (Fig 6A; p = 0.0229). As we had demonstrated that the immune evasive clones were also more immunotherapy resistant, we examined the effect of adding anti-PD1 immunotherapy to the T-cells *in vitro*. The addition of anti-PD1 significantly increased anti-cancer T-cell responses to the control clones and the IE1 clones, while T cell response to the IE2 clone was unchanged with the addition of anti-PD1 (p=0.36; Fig 6B).

**Figure 6.**
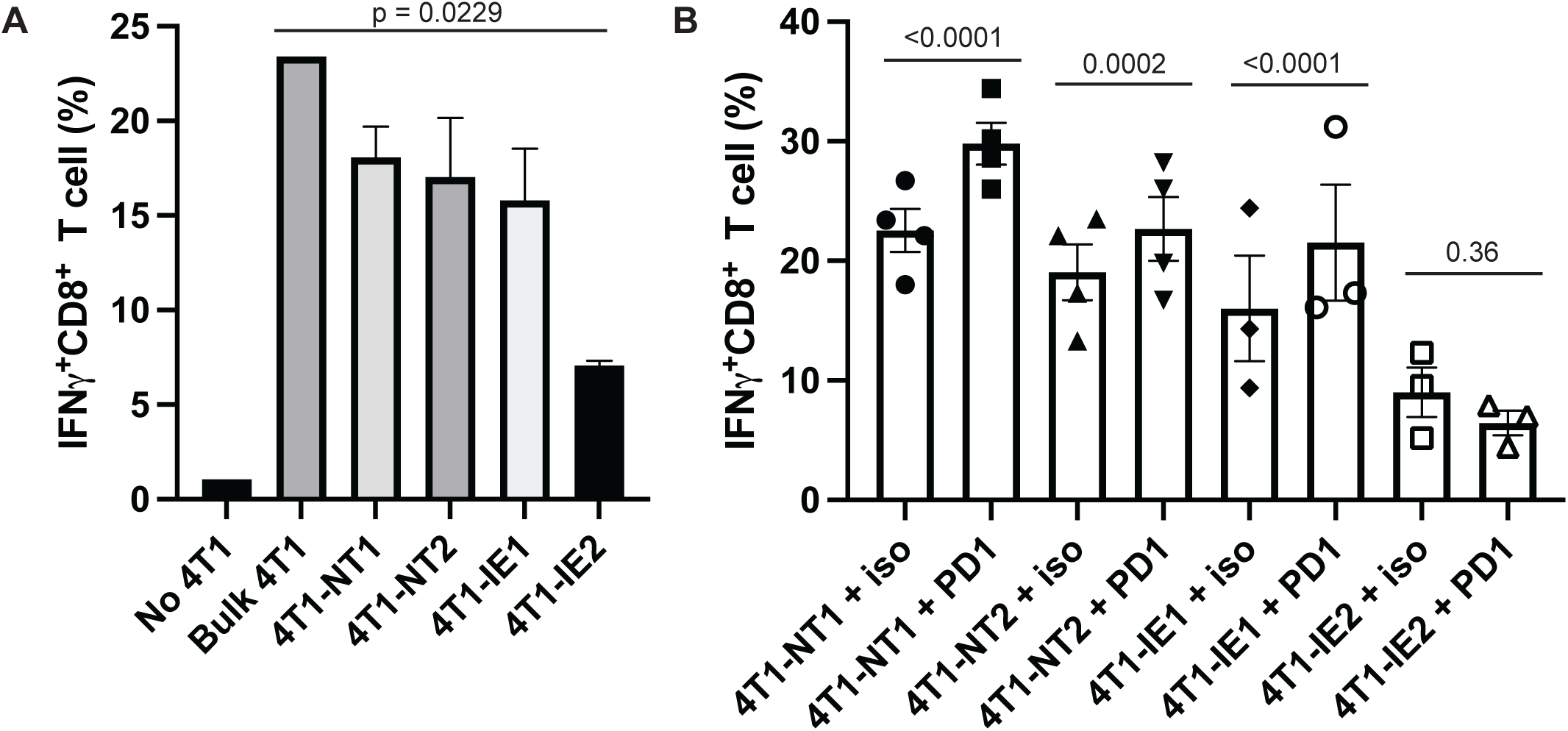
The 4T1-IE2 clone can directly suppress cytotoxic T-cell activation and prevent further T-cell activation by anti-PD1 immunotherapy. The 4T1-IE2 clone can directly suppress cytotoxic T-cell activation and prevent further T-cell activation by anti-PD1 immunotherapy. A. Measurent of intracellular IFNγ production by 4T1 specific CD8+ T-cells following co-culture with no 4T1 cells (No 4T1), bulk 4T1 cells (Bulk 4T1) and 4T1 clonal populations (4T1-N1; 4T1-NT2; 4T1-IE1; 4T1-IE2). For clonal populations n=3-4; one-way ANOVA with Dunnett’s post-hoc test. B. Measurent of intracellular IFNγ production by 4T1 specific CD8+ T-cells following co-culture with clonal 4T1 populations with or without anti-PD1 (PD1) or an isotype control (iso). n=3-4; FWER calculated using flow cytometry proportions fitted to a GLM with Bonferroni post-hoc correction.

This assay demonstrated that the IE2 clone uniquely evades activation of anti-cancer CD8 T-cells, and that the addition of anti-PD1 could not overcome the T cell suppression in response to this 4T1 clone. In contrast the IE1 clone induced a similar response to the control clones suggesting that it does not directly suppress CD8 T-cell activation.

### Overlapping gene signature is associated with poor survival in breast cancer patients

We had noted that the GSEA analysis showed some overlap in enriched gene-sets between IE1 and IE2. We thus reasoned that, in addition to having their unique immune evasion features, these clones may have some pathways in common. To identify immune evasion pathways common to both immunotherapy resistant clones, we identified overlapping gene expression changes (Sup Table 8). This analysis demonstrated that immunotherapy resistant clones had more genes with expression changes in common than either one had with either non-target clones (Fig 7A). We generated a heatmap of the top 50 upregulated and downregulated genes across all samples (Fig 7B) and performed GSEA analysis using C2 on the longer list (Sup Fig 13A, Sup Table 9). Only two gene sets had significant p values when multiple testing was considered, these were the HOXA5 gene set mentioned previously and a COVID-19 related gene set. Although not significant, there were several additional COVID-19 related gene sets from the same recent publication identified in the overlapping upregulated gene list suggesting an immune related role of these genes (*39*). We also identified three gene sets that were closely related and had clear associations with epigenetic regulation, “BENPORATH_ES_WITH_H2K27ME3”, “BENPORATH_PRC2_TARGETS” and “BENPORATH_SUZ_12_TARGETS”. In patient data, Ben-Porath and colleagues found negative enrichment of these genesets to be correlated with a stem-like phenotype and to be associated with poor prognosis (*40*). Further investigation showed the top genes driving these signatures were also negatively enriched in our derived gene signature, namely HHIP, CWH43, WNT10B, ABC3, CHN2 and CRIP1. This suggests a possible role for PRC2-mediated MHC-I suppression in our subclones.

**Figure 7.**
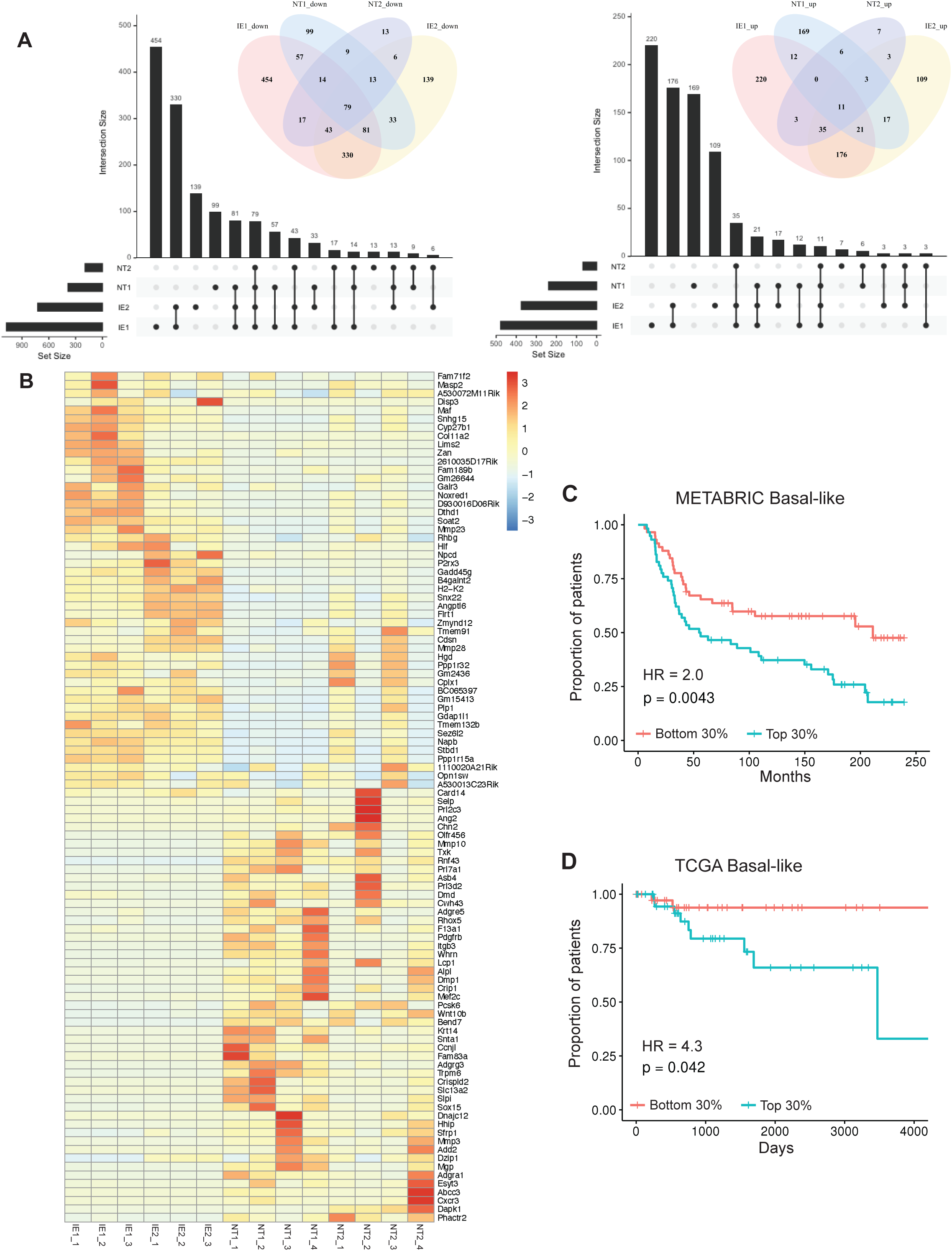
Overlapping gene signatures of the immunotherapy resistant clones show prognostic significance in basal-like breast cancer patients. A. Upset plots and Venn diagrams showing the overlap in significantly up-regulated (right) and down-regulated (left) genes between the two immunotherapy enriched clones (IE1 and IE2) and two control clones (NT1 and NT2). B. Heatmap of the top 50 overlapping up regulated and down regulated genes between IE1 and IE2 clones across all clonal cell lines. C. Kaplan-Meier analysis of overall survival of basal-like breast cancer patients in the METABRIC cohort based on whose breast cancer having the top 30% or the bottom 30% expression of the overlapping upregulated 25 gene signature. D. Kaplan-Meier analysis of overall survival of basal-like breast cancer patients in the TCGA cohort based on whose tumour having the top 30% or the bottom 30% expression of the overlapping upregulated 25 gene signature. In C and D, the Cox proportional hazards model was used to compute Hazard Ratios. Significance between stratification groups were computed using log-rank test statistics.

To understand the role of these genes in patients, we generated signatures from the top 25 upregulated and downregulated genes that had human orthologs and were detectable in both the METABRIC (*41*) and TCGA (*42*)datasets. We then analysed the association of these signatures with the survival of patients with basal-like breast cancer from these cohorts. We limited our analysis to basal-like breast cancer as we considered this the most relevant patient subgroup as the gene signature was derived from the 4T1 mouse model of basal-like breast. Although patients in these cohorts were not treated with immunotherapy, it has previously been demonstrated that immune features such as the number of tumour infiltrating lymphocytes or regulatory T cells influence prognosis in basal-like breast cancer patients (*43*). When we analysed overall survival, we observed that the upregulated gene signature associated with significant poorer outcome in both cohorts (METABRIC: p=0.0043, HR=2.0, Fig 7C; TCGA: p=0.042, HR=4.3, Fig 7D). In contrast, the downregulated gene signature did not show any significance in either cohort (data not shown). We generated heatmaps with unsupervised clustering to determine whether specific individual genes or groups of genes from the signature were driving the association with survival (Sup Fig 13B). In the TCGA dataset, we observed a number of clusters that seemed to associate more with survival, one of these included *FAM71F2, MASP2, HLF, PPP1R15A, MMP23B* and *LIMS2*. Interestingly, GADD34, encoded by *PPP1R15A*, had previously been demonstrated to be critical in blocking immunogenic cell death following chemotherapy, when it was inhibited, the chemotherapy response was improved in immunocompetent mice but not in immunocompromised mice (*44*). A second group of genes included *SEZ6L2*, which had been associated with survival in a number of cancers but not through an immune related mechanism (*45, 46*) There was no enrichment of proliferation or invasion gene sets in our GSEA analysis, suggesting that these processes were not behind the poor outcome of patients whose breast cancer highly expressed genes in the common upregulated signature.

Previous studies have shown that cytotoxic T lymphocytes (CTL) infiltration correlates with survival in basal-like breast cancer, so it is possible that our signature was a surrogate measure of T cell infiltration. To test this, we performed TIDE analysis (*47*) on TCGA and METABRIC cohorts followed by correlation analysis between the CTL signature score and our upregulated immune evasion signature score. This showed no correlation in the METABRIC cohort (Sup Fig 13C) and only a weak negative correlation in the TCGA cohort (Sup Fig 13D), suggesting little overlap between these two predictors of patient survival. Thus, it suggests we have discovered a novel immune related signature associated with patient survival in basal-like breast cancer. Future studies will be necessary to determine how the genes in this signature regulate survival, influence immune evasion and immunotherapy response.

## Discussion

Immunotherapy has revolutionised cancer therapy, with long-term responses seen in certain patient groups with a few types of cancer. Unfortunately, other patients having these same types of cancer and most patients having other types of malignancies, including breast cancer, have limited to no response to the current immunotherapies. Thus, our understanding of immune evasion in non-responsive cancer types needs significant improvement. We have addressed this by examining immune evasion at a clonal level and used this information to identify pathways that could be targeted to overcome immunotherapy resistance.

Here we show, for the first time in a solid tumour, that clonal immunoediting occurs and is enhanced by immunotherapy. Using the more immunotherapy sensitive EMT6 model we demonstrate that immunoediting occurs during the development of primary tumours and that immunotherapy leads to strong clonal selection. Most of the more immunotherapy resistant 4T1 cells are able to evade the immune system during the development of primary tumours but only a subset of them are able to evade the immune system during metastasis. This indicates that immune evasion is not a static process but requires ongoing regulation through tumour progression, even in a highly aggressive allograft model. These findings broadly agree with findings from a recent comprehensive genomic analysis of patient samples assessed across metastatic sites and over time (*3, 48*). These studies tracked clonal populations in metastatic lesions using whole genome sequencing, examined a number of immune correlates and identified immunoediting that was associated with the immune response. However, they were unable to examine clonal heterogeneity driven by epigenetic or transcriptomic changes and were limited in identification of rare clones by sequencing depth. Future clinical studies utilising single cell approaches to analyse multiple biopsies from individual patients over a time-course of treatment will be necessary to confirm the key findings of this study in patients. While extremely challenging, these studies are becoming more feasible with recent technological improvements.

Intriguingly the 4T1 model, unlike the EMT6 model, showed little immunoediting in the primary tumour. This suggests that either the vast majority of 4T1 cells are inherently resistant to immune control at the orthotopic site, or that 4T1 cells very rapidly set up a suppressive immune microenvironment that protects the majority of clones from immune mediated killing. The ability of 4T1 cells to induce myeloid derived suppressor cells could well contribute to a suppressive immune microenvironment, however, further studies would be necessary to further clarify this (*49*).

As we wanted to understand the role of immunotherapy in controlling metastatic disease we examined lung metastases from the 4T1 model following resection of the primary tumour. Lung metastasis occurs early in the 4T1 model with our studies indicating metastases can form following resection as early as day 13 (data not shown), and others showing the related 4T1.2 model robustly metastasizes by day 10 (*50*). This indicates that when adjuvant immunotherapy was given, these therapies were activating immune cells to target micrometastases that had already formed within the lungs. We postulate that while possible it is unlikely that circulating cancer cells were a major target of adjuvant immunotherapy as previous studies have indicated that circulating breast cancer cells only have a short half-life of 1-2.4 hours in circulation (*51*).

A number of previous barcoding studies in breast cancer had focused on metastasis and response to chemotherapy. However, these were performed in immunocompromised mice so that the role of the immune system in these processes was not addressed (*33, 52–54*). In the absence of a fully intact immune system it was demonstrated that specific clones have greater metastatic ability (*33, 52, 54*). Our results suggest that in the context of an intact immune system, a subset of these highly metastatic clones identified in these studies may have been recognised and removed via immunoediting. Similar to our results, these studies found that the dominant clone within the primary tumour generally was not the dominant clone in the metastases (*33, 52, 54*). Some of these studies also showed that chemotherapy treatment of PDX models led to a decrease in clonal abundance and diversity in relapsed disease, which is similar to what we found with immunotherapy (*53, 54*). Future studies combining immunotherapy with chemotherapy utilising a similar regimen to the atezolizumab plus nab-paclictaxel of the Impassion130 trial would give important insights into how combining these two treatment modalities affected clonal diversity following relapse (*55*).

Our and others’ studies have indicated that immunoediting can select for clones with immune evasive phenotypes irrespective of specific neoantigens (*2, 6, 56*). One previous study examining immunoediting at a clonal level used a fluorescent barcoding approach in a B cell leukemia model (*13*). While our findings broadly agree with the findings of that study, the study by Milo and colleagues was limited to five unique fluorescent clones that could be tracked and was confounded by the variable immunogenicity of these fluorescent proteins. DNA barcoding, in contrast, allowed for the labelling of thousands of clones and a much more precise identification of immune evasive clones. We could then isolate these clones and identify both common and variable features of immune evasion in them. Furthermore, this technique, unlike a fluorescent barcode approach, allowed us to demonstrate that clones that had greater immunotherapy resistance pre-existed in both EMT6 and 4T1 models, as these clones were enriched from the same starting pool of cells across replicate mice.

We identified and isolated two immunotherapy resistant clones from the 4T1 model. In depth analysis of these resistant clones demonstrated that they expressed genes of key immune evasive pathways (MHC-I and PD-L1) differently. Intratumoural heterogeneity (ITH) has previously been associated with resistance to immunotherapy in melanoma and lung cancer, with higher ITH being associated with resistance to immunotherapy (*26, 27, 57*). McGranahan and colleagues postulated that this was due to improved T cell killing of tumours with clonal neo-antigens. A non-mutually exclusive explanation is that clonal tumours are less likely to contain cancer cells with a pre-existing resistance mechanism to immunotherapy. These findings refine the concept of cancer immunoediting, demonstrating that there are clonal populations of cancer cells with variable resistance to the immune system. Based on their phenotype, these clones are either enriched or depleted by an active immune system and immunotherapy.

We identified a core overlapping gene expression profile between the two immunotherapy resistant clones. The common upregulated gene signature was able to stratify basal-like breast cancer patients into good and poor prognosis. This gene signature appeared to represent a novel immune evasion pathway associated with poor prognosis. Aside from *Ppp1r15a*, this signature did not contain genes known to be associated with immune evasion and it did not contain genes associated with other poor prognostic signatures, such as proliferation or invasion.

Because both T cells and NK cells are present during immunoediting in our models, this common signature likely enables cancer cells to evade both T cells and NK cells. Interestingly, our common signature does not strongly correlate with CTL infiltration, which indicates that this signature is not a surrogate for the lack of T cell infiltration and suggests that these genes likely do not regulate immune evasion by influencing immune cell recruitment. These common genes may offer new insights into developing therapeutic approaches to improve immunotherapy response in breast cancer. One of the common immunotherapy resistance genes we identified was *PPP1R15A*, which is consistent with its known role in immunogenic cell death in the response to chemotherapy (*44*). Recent studies in a mouse model of multiple sclerosis demonstrated that this pathway can be targeted using Sephin-1, a small molecule (*58*). Future studies are necessary to examine whether this compound or others targeting this pathway could synergise with immunotherapy or immunogenic chemotherapy to treat breast or other types of cancer. As breast cancers have relatively low mutational burden, it is likely that epigenetic factors may play a greater role than mutational events in driving ITH in breast cancer. One current approach to improve immunotherapy response under investigation is combining immunotherapy with epigenetic targeting drugs such as decitabine and HDAC inhibitors (*36, 59*). This combination has been shown to increase MHC protein expression and improve response to immunotherapy. However, epigenetic drugs may reduce the diversity of clones and overcome other epigenetically driven immune evasion mechanisms in addition to enhancing MHC expression (*60*). Further research is needed to test this hypothesis more fully. Our results, however, suggest that while demethylating agents could increase MHC-I expression in the MHC-I low immunotherapy resistant clone, they did not increase it above the baseline seen in the parental 4T1 cell line. This is corroborated by a recent study demonstrating that while treating breast cancer patients with demethylating agents could increase MHC-I expression in most patients’ tumors, a subset appeared resistant to this therapy (*36*). This suggests that while epigenetic treatments may improve the proportion of patients responding to immunotherapy, in some cases pre-existing clones could still mediate resistance to this combination.

A limitation of this study is the reliance on mouse cell line models, which do not recapitulate early stages of tumorigenesis and do not represent the full diversity of human breast cancer. However, syngeneic allograft models have delivered central insights about the immune response to cancer and demonstrated the utility of immunotherapies (*61*). Another limitation is that the integration of the barcode and selection markers into the genome and the potential immunogenicity of red fluorescent protein (RFP) could affect the phenotype of these cancer cells. We and others have found in previous studies that some fluorophores and luciferase were immunogenic and negatively affect tumour growth and metastasis in the 4T1 model (*29, 62*). However, we found that tumour growth and metastasis were unaffected by RFP expression in this study. While the introduction of DNA barcodes could have influenced the phenotype of specific clones, we feel that this is unlikely given that no dramatic impact on the expression of the genes closest to the integration site. Furthermore, none of the genes associated with integration sites was identified to be significantly involved in cancer cell evasion of CD8 T cell responses in a recent CRISPR screen .

To survive in any given system, cancer cells must utilise a number of mechanisms to avoid more than just immune destruction, as extensively reviewed in the recent work of Hanahan (*64*). Not only are IE1 and IE2 immune evasive, they are also highly metastatic and by their very nature must be able to grow independent of anchorage, as well as possessing abilities to engraft in both the mammary fat pad and lungs, shed from the primary tumour prior to resection and resist all other mechanisms of host anti-cancer response. We attribute the decrease in unique clones present in the lungs of control treated 4T1 tumour bearing mice compared to the primary tumour to precisely this; only some clones within the engrafted tumour possess the required abilities to be able to successfully metastasise. By making comparisons between controlled conditions, e.g. immunotherapy treated lungs vs control treated lungs, we believe we have effectively demonstrated immunoediting occurring *in vivo*.

Overall, this study has demonstrated that immunoediting occurs at the clonal level in primary tumours and that a second round of immunoediting occurs during metastasis. Immunotherapies dramatically enhanced immunoediting, but pre-existing resistant populations were still responsible for relapse. The large reduction in clonal diversity following immunotherapy in the 4T1 model, which is known to be poorly responsive to immunotherapy, suggests that slight improvements through combination therapy could eliminate the remaining clones and lead to dramatic improvements in survival. By isolating immunotherapy resistant clones and characterizing them, we identified common and distinct immune evasion pathways. We anticipate that through targeting pathways identified in this study, in particular common pathways, it will be possible to further reduce the numbers of resistant clones and improve the efficacy of immunotherapies.

## Acknowledgments

This work is supported by research grants from The National Breast Cancer Foundation (NBCF) of Australia (IN-17-02; IIRS-19-106) and supported by the generosity of John McMurtrie (AM) and Deborah McMurtrie and The Petre Foundation. LAB is the recipient of an Australian Government research training (RTP) scholarship. AS is the recipient of a Senior Research Fellowship from the National Health and Medical Research Council of Australia. SJ is supported by a research fellowship from the NBCF. The authors would like to acknowledge Dr Tim Peters and thank him for his valuable statistical advice. We acknowledge the contribution of Isabelle Shapiro and Cheryl Grant as consumer advocates. This manuscript was edited at Life Science Editors.

## Author Contributions

L.A.B. performed experiments, analysed and interpreted the data, and drafted the manuscript. N.B. performed bioinformatic analysis of the barcoding and RNAseq data. J.Y. performed experiments. S.Z.W. performed bioinformatic analysis of patient cohorts. N.D. performed bioinformatic analysis of the DNAseq data. D.L.R. performed bioinformatic analysis of the barcoding data. C-L.C. performed experiments. G.A. analysed and interpreted the data. D.J.Z and B.S.P performed experiments and analysed data. A.S. helped conceive the study, analysed and interpreted the data, and helped draft the manuscript. S.J. conceived the study, performed experiments, analysed and interpreted the data, and drafted the manuscript.

## Methods

### Cells

4T1 cells were obtained from ATCC. 4T1 cells were grown in RPMI (Gibco) supplemented with 10% FCS, D-Glucose, Sodium Pyruvate, 2mM HEPES, and Penicillin/Streptomycin. EMT6 cells were obtained from ATCC. EMT6 cells were grown in Waymouth’s MB 752/1 Medium supplemented with 15% FCS and 2mM L-glutamine. Cells were tested for mycoplasma contamination using the MycoAlert Mycoplasma Detection Kit, 100 Test Kit (Catalog# LT07-318).

### Cellular DNA barcoding

The ClonTracer library was a gift from Dr Frank Stegmeier (Addgene #67267). Lentiviral particles containing the high-complexity barcode library were produced by transfecting 293T cells. 4T1 and EMT6 cancer cell lines were barcoded by lentiviral infection using 0.8 µg/ml polybrene. Cells from each line were infected with a target MOI of 0.1, corresponding to 10% infectivity to ensure single lentiviral integration. Cells that received a barcode were then sorted based on the RFP reporter protein using a BD FACSAriaII, these cells were then expanded and frozen into a number of aliquots for the subsequent experiments. 4T1 cells were generated with two different barcode complexities, one with ∼5000 barcodes (4T1 BC5000) and one with ∼300 000 barcodes. The EMT6 cells and the high complexity 4T1 cells were passaged twice following cell sorting, frozen and cells from these aliquots were thawed and passaged once more prior to transplantation. The low complexity 4T1 cells (4T1 BC5000) were a subpool derived from the higher complexity 4T1 cell line, these were passaged an additional 3 times to expand and freeze and were then thawed and used as described above.

### *In vivo* experiments

All animal experiments were approved by the Garvan Institute of Medical Research/St. Vincent’s Hospital Animal Experimentation Ethics Committee.

Immunocompetent BALB/c mice and immunocompromised NOD.Cg-*Prkdc^scid^ Il2rg^tm1Wjl^*/SzJ (NSG) mice aged 6-to-8 weeks were obtained from Australian BioResources (Moss Vale, Australia) and housed at the Garvan Institute of Medical Research.

### *In vivo* tumour growth

For tumour transplantation, barcoded EMT6 cells (ATCC, USA) were resuspended in Matrigel 2.5x10^5^ cells in 100 ml volume were injected into the 4^th^ inguinal mammary gland; barcoded 4T1 cells (ATCC, USA) were resuspended in PBS and 5x10^4^ cells in a 10 ml volume were injected into the 4^th^ inguinal mammary fat pad. For studies with the 4T1 model primary tumours were surgically resected at day 15. At resection or ethical endpoint tumours and whole lung tissue were removed, minced and snap frozen in liquid nitrogen for barcode analysis.

### Immunotherapy treatment

Mice were treated with four 200 μg doses of either combination immunotherapy antibodies via intraperitoneal injection: anti-CTLA4 (BE0032), anti-PD1 (BE0146), or isotype control antibodies Armenian hamster IgG (BE0091), Rat IgG (BE0089) all from BioXCell (Lebanon, NH, USA). Antibodies were given every 2-3 days from day 10 after tumour implantation for the EMT6 model and following resection of primary tumour on day 15 for the 4T1 model.

### CD8 T cell, CD4 T cell and NK cell depletion

Starting one day prior to primary tumour resection mice were given 100 μg of depleting antibodies for CD8 T cells (anti-CD8; BEO223; BioXCell), or NK cells (anti-Asialo-GM1; 986-10001; Novachem), or CD4 T cells (anti-CD4; BE0003-1; BioXCell) or isotype control antibodies. Antibodies were then given every 2-3 days for a total of 4 doses.

### Isolation of buffy coat and flow cytometry to confirm depletion of key cell types

Whole blood was collected into K2EDTA coated tubes (BD, cat# 365973) from mice two days after the last intraperitoneal injection of depleting or control antibodies. Buffy coat was isolated by spinning blood at 800g at room temperature for 10 minutes and removing the small layer of the buffy coat. Any red blood cells were lysed using Pharmlyse (BD, 5 min) and quenching with FACS buffer (DPBS supplemented with 2% FCS and 2% HEPES). Cells were washed and stained with BV711 conjugated anti-mouse CD8a (Biolegend, 1:200, cat# 563046), PE conjugated anti-mouse CD4 (Biolegend, 1:200, cat# 116006), and APC conjugated anti-mouse NKp46 (Biolegend, 1:200, cat# 137607). Cells were washed 3 times before staining with DAPI. Data was collected using the BD LSR Fortessa (FACSDiva 8.0.1) .

### DNA extraction

Frozen tissue samples were lysed in 5 ml QIAGEN buffer P1 (with RNaseA) and 0.5% SDS within a Miltenyi M-Tube (# 130-096-335). Samples were processed on the gentleMACS or gentleMACS Octo using the RNA_02 program. DNA was then extracted using a standard phenol/chloroform process.

### Targeted Barcode PCR and Sequencing

All samples underwent targeted barcode PCR amplification according to the updated version of the original protocol (*30*) available on the Addgene website (https://www.addgene.org/pooled-library/clontracer/). Specific PCR products (180 bp) were gel purified, quantified by Qubit 2.0 fluorometer (ThermoFisher Scientific, Waltham, MA USA) and pooled into a library. Prior to sequencing, an equal combination of additional PCR products containing two inverse barcodes (GACTCAGTGTCAGACTGAGTGTCTGACTGT and CTGAGTCACAGTCTGACTCACAGACTGACA) plus the PhiX Control V3 (Cat. FC-110-3001, Illumina, CA, USA) were spiked in to balance the nucleotide distribution within the library. Samples were sequenced using a custom sequencing primer (GCGACCACCGAGATCTACACACTGACTGCAGTCTGAGTCTGACAG) and the NextSeq^®^ 500/550 Mid Output Kit v2 - 150 cycles (FC-404-2001, Illumina, CA, USA) on the Illumina NextSeq^®^ platform.

### Barcode Analysis

Barcode composition analysis and calculation of barcode overlap between samples was performed as indicated in the original protocol (*30*) and updated Python scripts available from the Addgene website (https://www.addgene.org/pooled-library/clontracer/).

Further analysis was performed using R statistical framework and packages EntropyExplorer for analysis of differential Shannon Entropy (*65*), DEBRA for differential barcode expression (*66*), and libraries fishplot and UpSetR for visualisation purposes. Pheatmap (v1.0.12) was used with default parameters. Clustering distances for rows and columns were euclidean and complete clustering linkage.

### Generating clonal cell lines

Cells of interest were isolated from the barcoded parental population using a sub-pooling approach.

The barcoded 4T1 BC5000 cells were seeded into a 96 well plate at a density of 150 cells per well. At approximately 80% confluence, cells were trypsinised and split identically into two plates. One plate was viably frozen in freezing media (10% DMSO, 40% FCS, 50% 4T1 media). DNA was extracted from one plate using the Promega SV Wizard Genomic DNA kit. Target barcodes of each sample were PCR amplified and sequenced using the method described above.

After sequencing, wells containing cells with the target barcodes were thawed, pooled and seeded at 40 cells/well in a 96 well plate. Media was changed every three days for 8 days before cells were split into two identical plates as above. One plate was viably frozen in freezing media, while DNA was extracted and prepared for targeted sequencing as above.

Wells with the highest proportion of target barcodes were revived into a 6 well plate and grown for 4 days before being single-cell sorted by BD FACSAria II into 96 well plate. Sorted single cells were grown in conditioned media for 5 days before being changed to 4T1 media and grown until 80% confluence. As previously, cells were lifted and split identically into two plates – one for freezing and one for targeted sequencing. Once wells containing single cells clones of the cells of interest were identified, target wells were revived. Cells were expanded before being aliquoted and viably frozen for future experiments.

Barcoded sequences of isolated cells were confirmed by targeted Sanger sequencing of barcode regions.

### Bulk RNA sequencing

RNA was extracted from established subclonal cell lines using the QIAGEN RNeasy Mini Kit. 3-4 unique clonal cell populations were sequenced for each barcode. Libraries were prepared using the KAPA RNA HyperPrep Kit with RiboErase, and sequenced on the NextSeq500 platform using a High Output V2.5 300 cycle kit.

### Transcriptome analysis

FastQ files from sequencing libraries were first trimmed with FASTQC v0.11 *Andrews S.* (*2010*)*. FastQC: a quality control tool for high throughput sequence data. Available online at:* http://www.bioinformatics.babraham.ac.uk/projects/fastqc. Raw reads were subsequently mapped to the mouse transcriptome (Gencode release M9, GRCm38.p4), to the mouse genome (mm10 assembly), with STAR aligner v.2.4.1d, allowing for multimapping reads (*67*). The reads were counted over gene models with RSEM, v.1.2.18 (*51*). Differentially expressed genes and repeat elements were defined with EdgeR with FDR<0.01 (*68*). EdgeR uses genewise negative binomial generalised linear models through the function glmQLFTest (*69*). Genes that with less than 10 reads across three samples per group were omitted from the analysis. The model involved comparison between IE samples, NT samples and the bulk 4T1 cell line.

### Survival analysis

To assess the clinical relevance of our isolated immune evasion clones, we assessed the association between the gene signatures derived from our bulk RNA-Sequencing studies with the overall survival of basal (PAM50) breast cancer patients from the METABRIC and The Cancer Genome Atlas (TCGA; https://www.cancer.gov/tcga) cohorts. Mouse gene signatures were first converted to human orthologs using the biomaRt package (*70*). Shared up-regulated genes across both immune evasion clones IE1 and IE2 were then filtered, and only genes detected in each expression cohort were considered. For each tumour from the bulk cohort, signature scores were computed based on the average expression of the top 25 genes ranked by log fold change. Patients were then stratified based on the signatures scores into the top 30%, middle 40% and bottom 30% groups. Survival curves were generated using the Kaplan Meier method with the ‘survival’ package in R (https://cran.r-project.org/package=survival). The Cox proportional hazards model was used to compute Hazard Ratios. We assessed the significance between groups using the log-rank test statistics.

### Gene set enrichment analysis

Gene set enrichment analysis (GSEA) was carried out using the GSEA desktop app (4.1.0) and DEGs generated as described above. GSEA was run using preranked list of significantly differentially expressed genes, ranked by log fold change. The molecular signatures database (MsigDB v7.5.1) hallmark and curated (C2) gene sets were used for analysis.

### Whole genome sequencing

DNA was extracted from established subclonal cell lines using the QIAGEN DNeasy blood & tissue kit. Libraries were prepared using the Roche KAPA PCR-free library preparation kit and genomes were sequenced on the HiSeq X platform to a depth of ∼30x.

### Whole genome analysis

Fastq files from the WGS were firstly aligned to mouse genome reference mm10. The output bam files were subsequently used for copy number analysis. Copy number analysis was performed using a R package cn.mops (*71*) in “paired mode” with a window length of 10kb. Reads were aligned to the BALB/c reference genome using BWA before being indexed and sorted with Novosort. Reads that mapped incompletely to the reference genome were then mapped to the barcode plasmid sequence with BWA and sorted and indexed with Novosort. Read pairs where only one pair mapped to the barcode plasmid sequence were blasted against mm10 to establish the barcode plasmid insertion site.

### Flow cytometry for MHC-1 and PD-L1

The 4T1 subclones (IE1, IE2, NT1, NT2), as well as the parental 4T1 bulk population were revived and passaged three times before being seeded into a 6 well plate at a density of 200 000 cells per well. At approximately 80% confluence, cells were collected into FACS buffer (DPBS supplemented with 2% FCS and 2% HEPEs) for flow cytometry. Cells were stained with a mastermix of APC conjugated anti-mouse CD274 (Biolegend) and Alexa Fluor488 conjugated anti-mouse H2-kD (Biolegend) at a final concentration of 1:200 for 20 minutes. Cells were washed three times with FACs buffer before being stained with DAPI and run on the BD FACSCanto II flow cytometer, utilising BD FACSDIVA software. Data was analysed in FlowJo (version 10.6.1) and median fluorescence intensity of live, single cells was calculated.

### Treating cells with 5-Aza-2’-deoxycytidine and flow cytometry for MHC-I

5-Aza-2’-deoxycytidine (5aza) was sourced from Sigma and reconstituted in DMSO according to manufacturer’s instructions. Subclones (IE1, IE2) and the parental 4T1 cell line were seeded into a 24 well plate at a density of 8000 cells per well in 4T1 media (previously described). Cells were allowed to settle overnight before being treated with 5aza at 200nM, 100nM or 50nM for 72 hours. 5aza was removed and cells were cultured in media only for 24 hours before being collected for flow analysis. Cells were stained with Alexa Fluor488 conjugated anti-mouse H2-kD (Biolegend) at a concentration of 1:200 in FACS buffer for 20 minutes. Cells were washed three times after staining before being stained with DAPI. Data was collected using the BD FACSCanto II flow cytometer with BD FACSDIVA software. The resulting data was analysed using FlowJo (version 10.6.1) and media fluorescence intensity of live, single cells was calculated.

### Treating cells with interferon gamma and flow cytometry for MHC-I

Active mouse interferon gamma (IFNγ) was sourced from Abcam and reconstituted in sterile water, as per manufacturer’s instructions. Subclones (IE1, IE2) and the parental 4T1 cell line were grown in a 24 well plate until approximately 70% confluence was achieved. Cells were then treated with IFNγ (100ng/ml) for 24 hours. Cells were stained with Alexa Fluor488 conjugated anti-mouse H2-kD (Biolegend) at a concentration of 1:200 in FACS buffer for 20 minutes. Cells were washed three times before being stained with DAPI. Data was generated using the BD FCSCanto II flow cytometer with BD FACSDIVA software. Analysis was carried out using FlowJo (version 10.6.1) and median fluorescence intensity of live single cells was calculated.

### Dissociation of metastatic lung tissue and flow cytometry for MHC-I and PD-L1

Lungs were minced with scissors and dissociated in GentleMACS C tubes (Miltenyi Biotec) with collagenase (1mg/ml, Type 1A from clostridium histolyticum, Sigma, cat # C9891) in collagenase buffer (RPMI-1640 supplemented with 2.5 % FBS and 1% HEPES), shaking, 37°C for 50 minutes. After shaking, samples were passed through 70um MACS SmartStrainers (Miltenyi Biotec) and washed with PBS, samples were treated with DNase (1mg/ml, from DNase 1 from bovine pancreas, Sigma, cat# CD25) for 3 minutes and quenched in FACS buffer (DPBS, Gibco, supplemented with 2% FBS and 2% 1M HEPES). Samples were washed twice in FACS before FC block with Mouse BD Fc Block (BD, cat# 553141) for 15 minutes. Cells were stained with Alexa Fluor488 conjugated anti-mouse H2-kD (clone SF1-1.1, Biolegend, cat# 116610) and APC anti-mouse CD274 (B7-H1, PD-L1; clone 10F.9G2; Biolegend; cat# 124311) at a concentration of 1:200 in FACS buffer for 20 minutes. Samples were washed three times before being stained with DAPI. Data was generated using the BD LSR Fortessa flow cytometer with BD FACSDIVA software. Analysis was carried out using FlowJo (version 10.6.1).

### Generation of poly-specific cytotoxic T lymphocyte lines

4T1-specific CD8+ T cell lines were derived as previously described (*37*) Briefly, Balb/c mice were immunised with 1x106 irradiated 4T1 cells via intraperitoneal injection. Spleens were harvested >30 days post-immunisation and re-stimulated in vitro with irradiated 4T1 cells and cultured in RPMI containing 10% FBS and 10 IU IL-2/mL for 14 days before experimental use.

### Functional assessment of tumor cell line immunogenicity

4T1 functional assessment was performed as previously described (*38*). Briefly, 4T1 cell line variants were cultured in the absence or presence of 100 ng/mL recombinant IFNγ in 37°C 5% CO2 for 24 h. 5 x 104 treated cells were washed twice with PBS and incubated with 5 x 103 4T1-specific CTL in the presence of 10 µg/mL Brefeldin A in 37°C 5% CO2 for 5 h. For anti-PD1 experiments: 4T1-specific CD8+ T cells were incubated in the presence of 10 µg/mL anti-PD1 (clone: RMP1-14, BioXcell) or isotype control (clone: 2A3, BioXcell) for 30 min, RT. Samples were then stained with cell surface antibodies for 20 min, 4°C (α-CD8, clone: 53-6.7, 1:300, eBioscience), washed with PBS, fixed with 1% paraformaldehyde (15 min, RT, dark), permeabilized and stained for intracellular proteins (α-IFNγ, clone: XMG1.2, 1:300, eBioscience) in the presence of 0.4% Saponin for 30 min, 4°C, and analysed on a FACSymphony A5 instrument (BD).

### Data availability

RNA-seq data have been deposited in the ArrayExpress database at EMBL-EBI (www.ebi.ac.uk/arrayexpress) under accession number E-MTAB-10027.

DNA-seq data have been deposited in the ArrayExpress database at EMBL-EBI (www.ebi.ac.uk/arrayexpress) under accession number E-MTAB-10027.

---

**Sup Figure 1.**
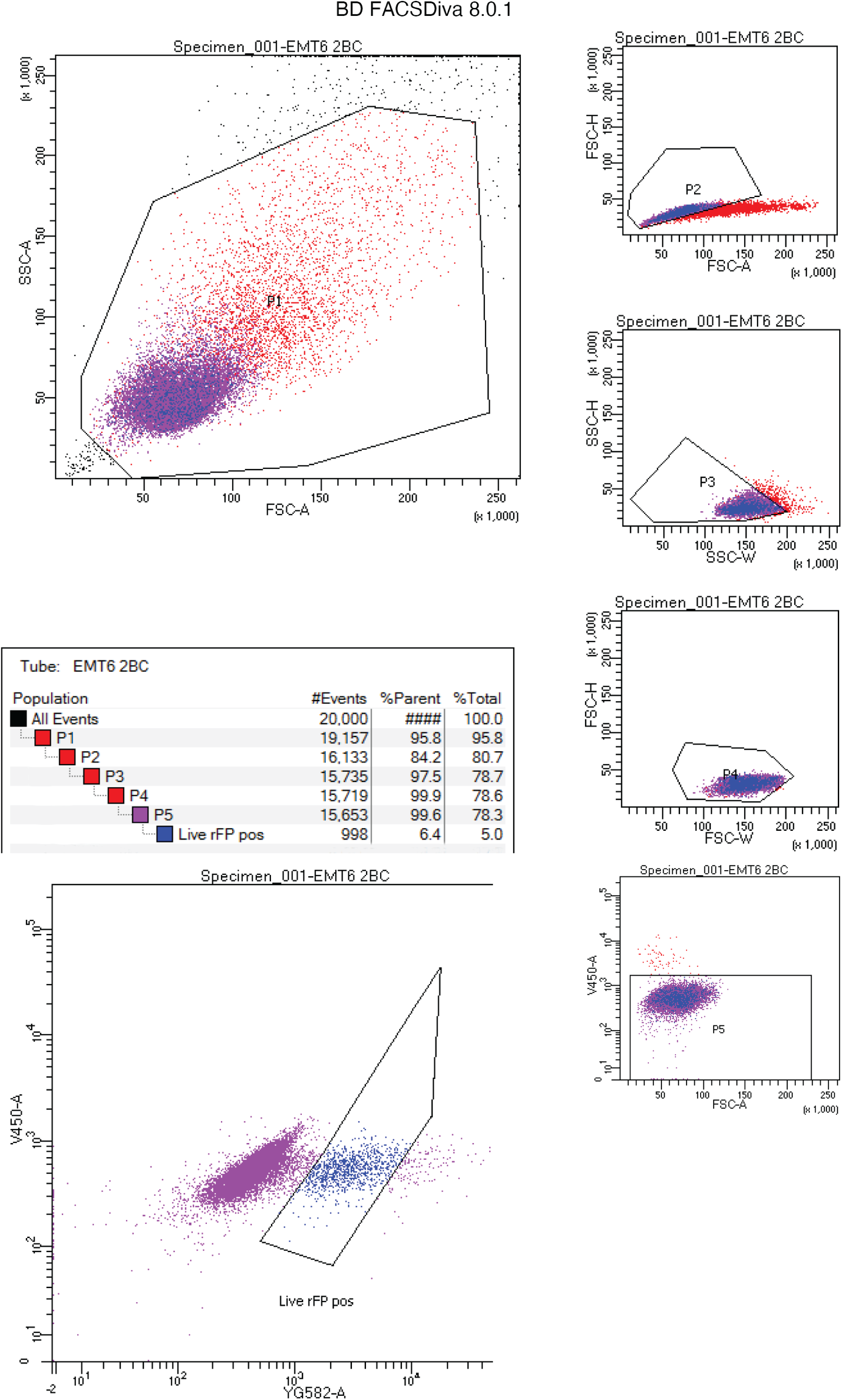
Gating strategy showing isolation and selection of barcode transfected Red Fluorescent Protein (RFP) positive EMT6 cells

**Sup Figure 2.**
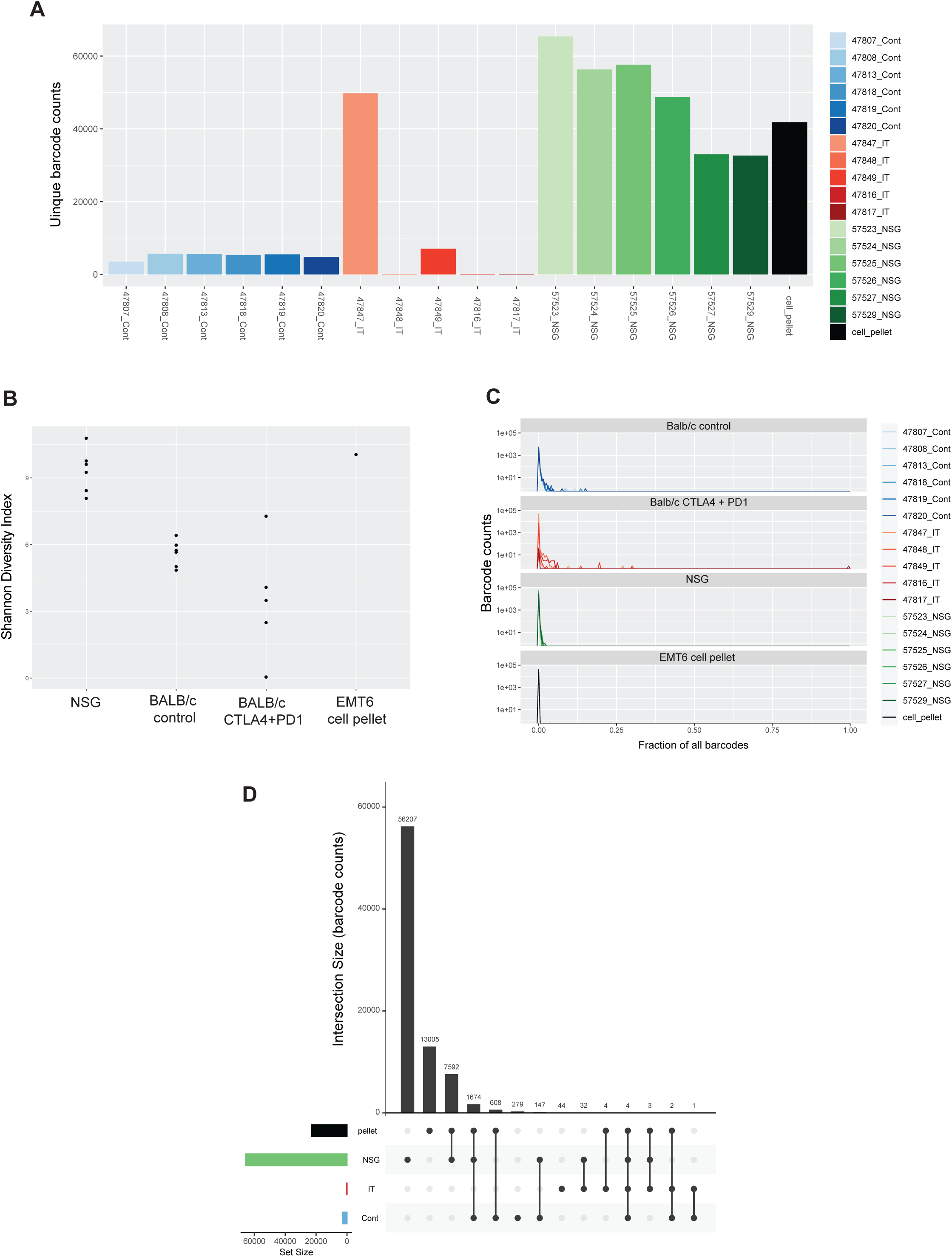
Unique barcode counts and barcode distributions across EMT6 primary tumours and matching cell pellets. A. Unique barcode counts from EMT6 tumour bearing mice. B. Shannon diversity index. C. Distribution plots of barcode counts from individual mice, grouped by strain and treatment group. D. Upset plot showing unique barcode counts and overlap between the EMT6cell pellet and tumour bearing NSG mice or immunocompetent immunotherapy treated (IT) or control treated (Cont) mice.

**Sup Figure 3.**
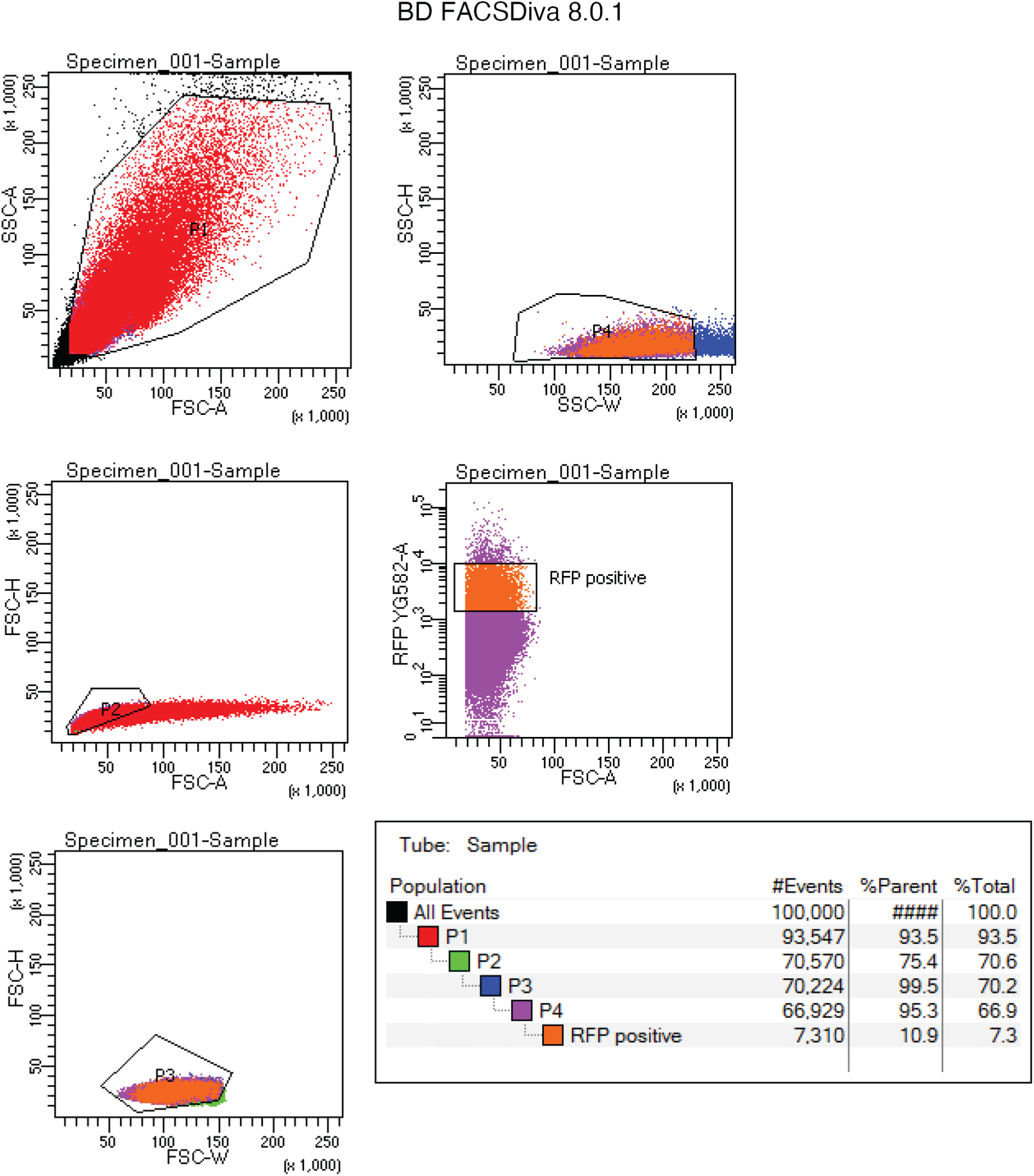
Gating strategy showing isolation and selection of barcode transfected Red Fluorescent Protein (RFP) positive 4T1 cells.

**Sup Figure 4.**
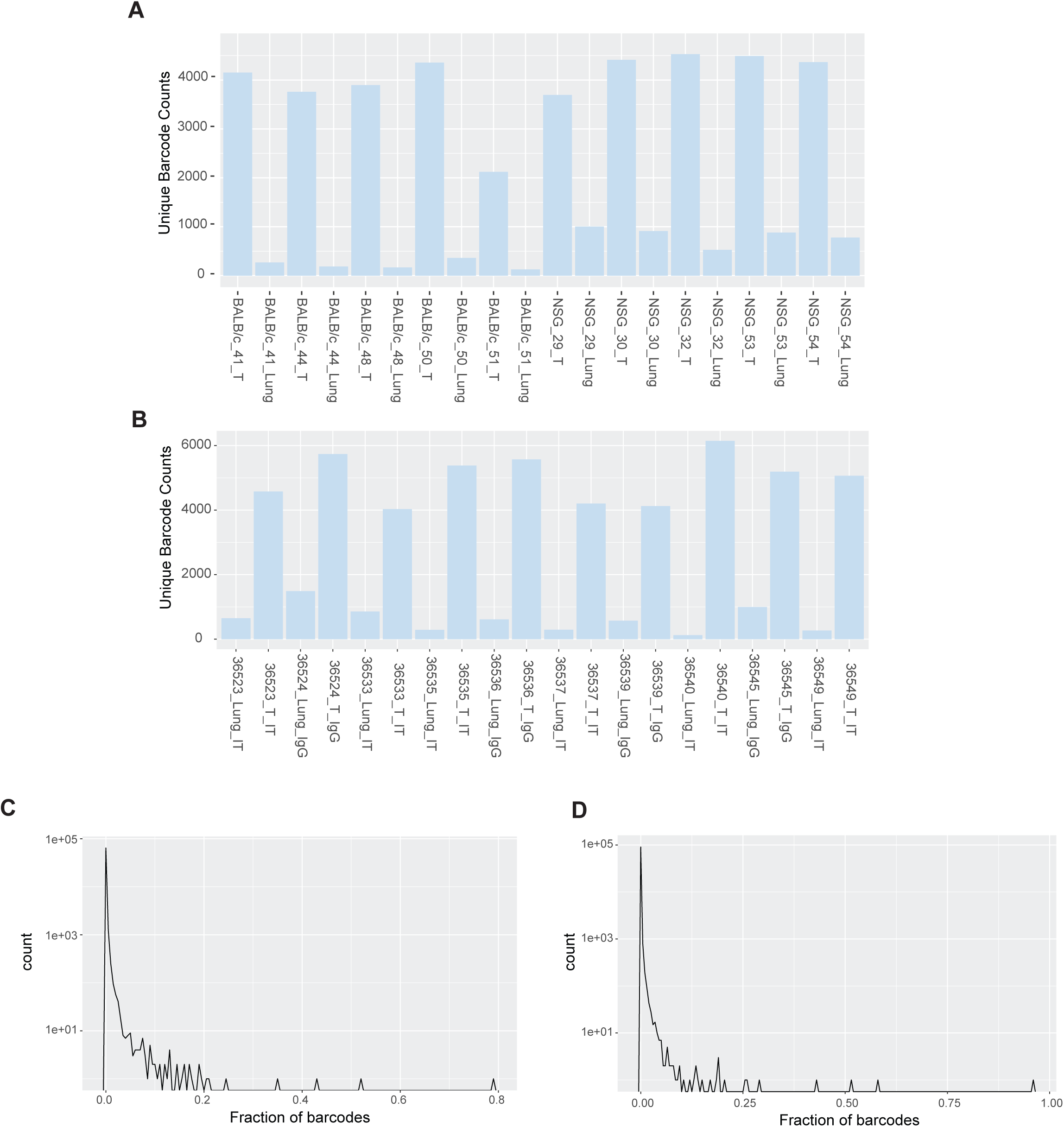
Unique barcode counts and barcode distributions across 4T1 primary tumours and matched lungs in NSG and Balb/c mice. A. Unique barcode counts detected in matched primary tumour (T) or lungs from 4T1 tumour bearing NSG or Balb/c mice. B. Unique barcode counts detected in matched primary tumour (T) and lungs from 4T1 tumour bearing mice treated with either combined immunotherapy (IT) or control antibodies (IgG) C. Distribution plot of barcode counts from all (NSG and Balb/c) mice. D. Distribution plot of barcode counts from Balb/c mice treated with combination immunotherapy or control antibodies.

**Sup Figure 5.**
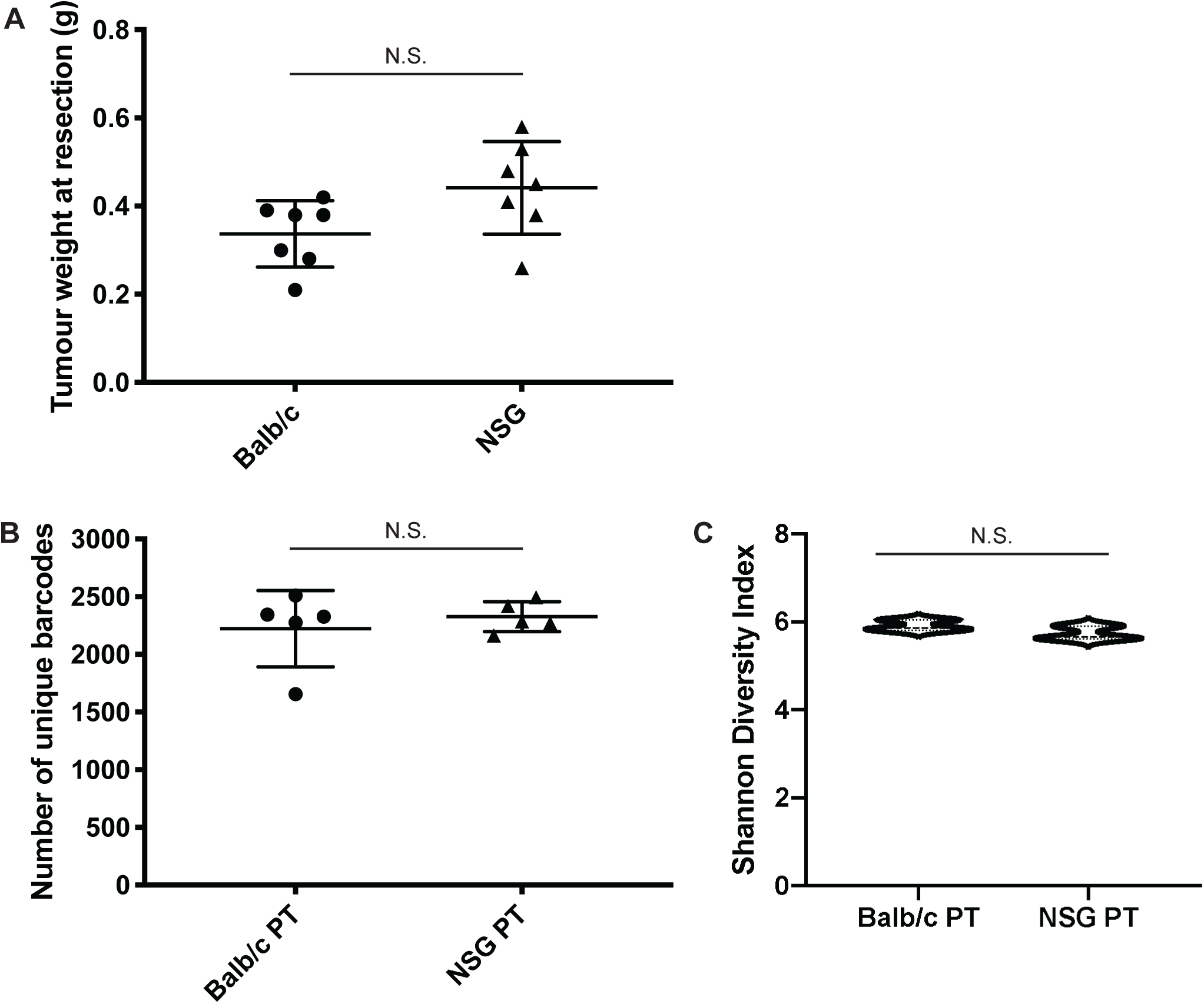
4T1 primary tumour growth and barcode diversity in primary tumours. A. Primary tumour mass. B. Total number of unique barcodes identified in primary tumours of Balb/c or NSG mice. C. Shannon diversity analysis comparing primary tumours from Balb/c and NSG mice.

**Sup Figure 6.**
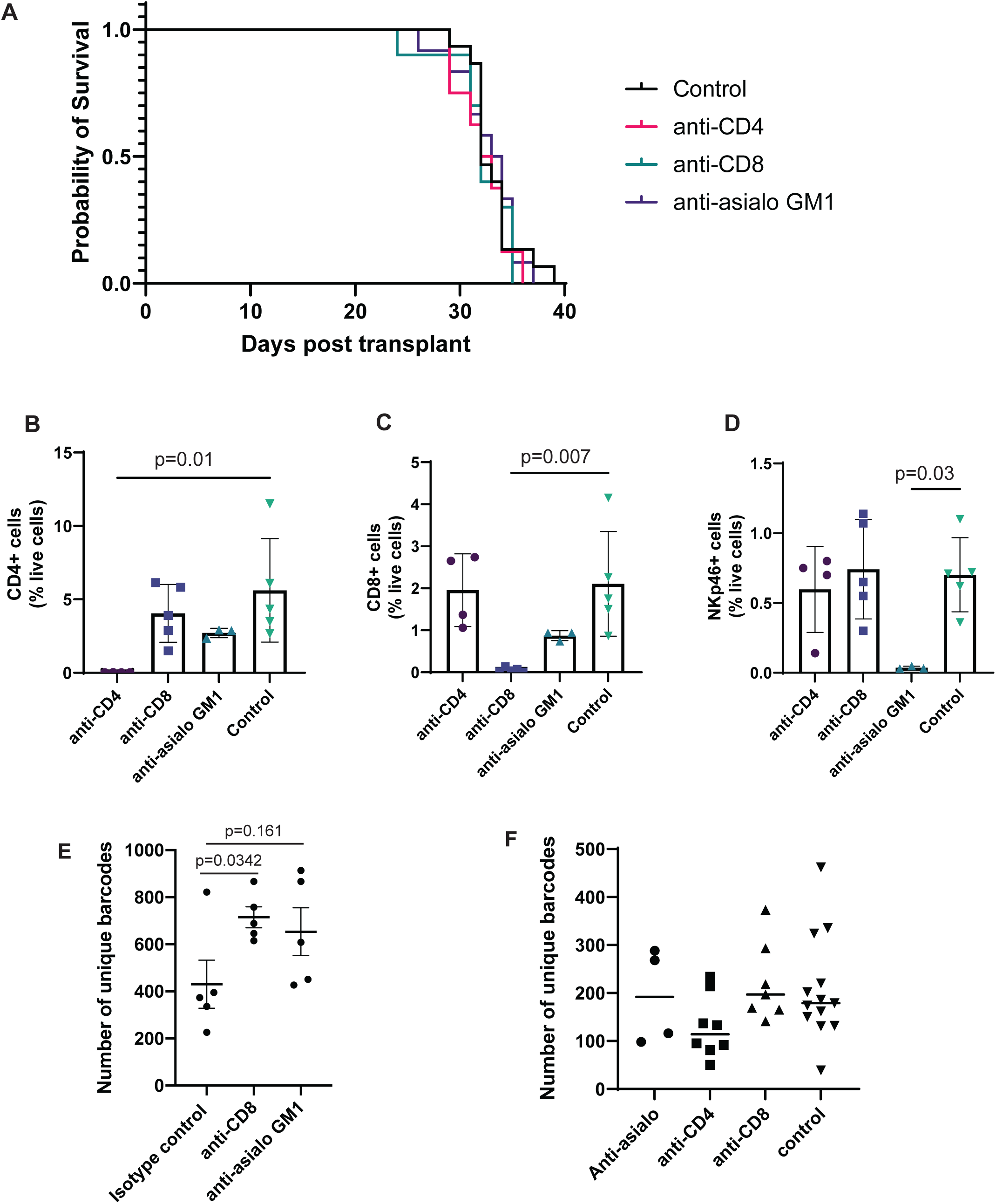
Depletion of CD8+ T cells with anti-CD8, CD4+ T cells with anti-CD4 or NK cells with anti-asialo GM1 does not significantly affect survival or barcode diveristy of mice bearing 4T1 tumours. A. Kaplan-Meier survival analysis of Balb/c mice transplanted with 4T1 cells, primary tumour was resected on day 15. Depletion of target cell types was initiated 1 day prior to resection, on day 14. 8-15 mice per treatment group. B-D. Flow cytometry to confirm depletion of target cell types. Buffy coat was collected on day 22, 2 days after the final dose of depleting antibodies was given. One way ANOVA with Tukey’s HSD for multiple comparisons. E. Initial experiments showed a small increase in unique barcode number in the lungs when mice were treated with CD8-depleting antibodies. F. Repeated experiments showed depletion of key cell types does not significantly change barcode number in the lungs at endpoint.

**Sup Figure 7.**
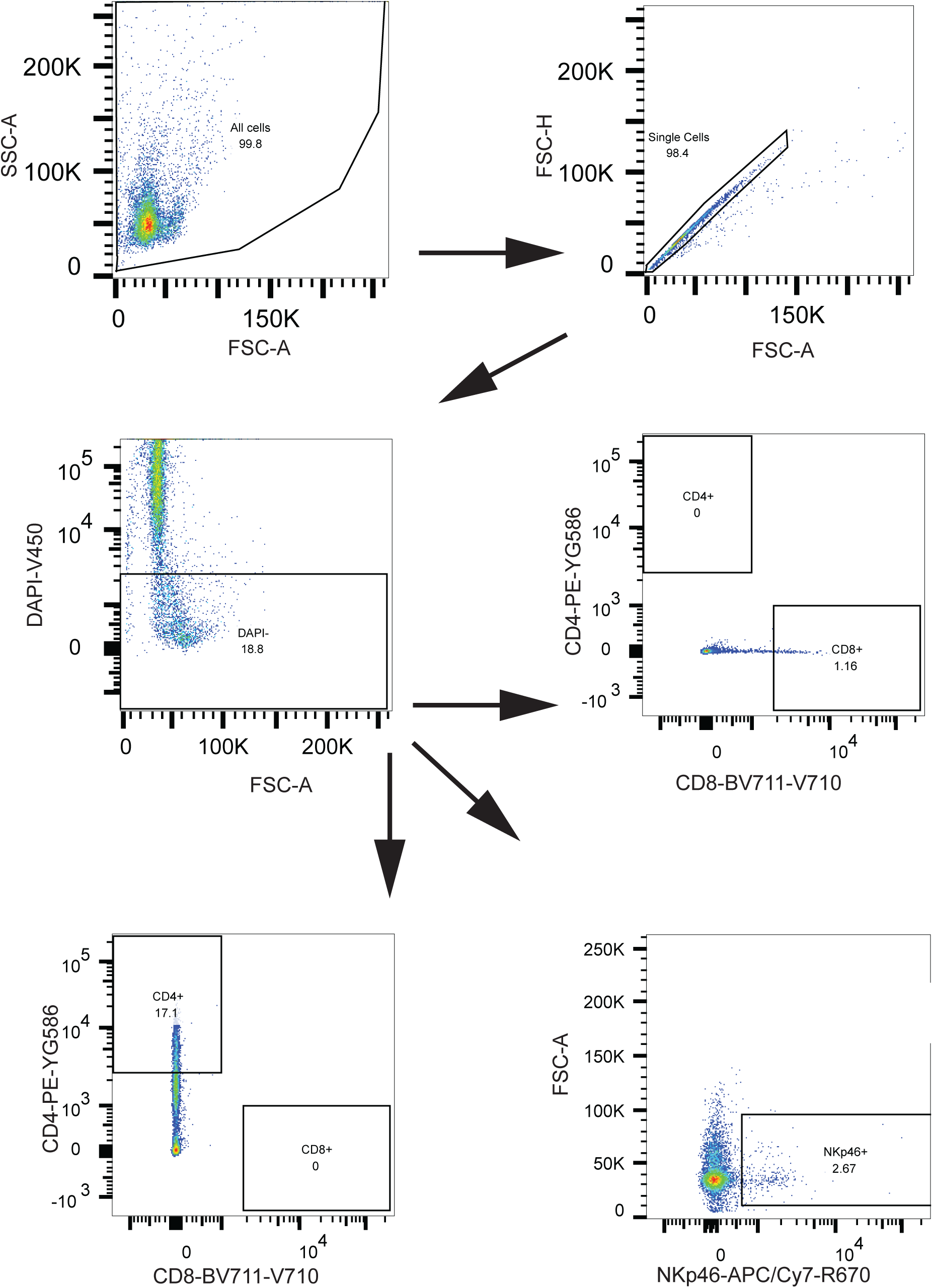
Gating strategy to confirm depletion of target cell types in 4T1 tumour bearing mice, treated with either anti-CD4, anti-CD8 or anti-asialo GM1.

**Sup Figure 8.**
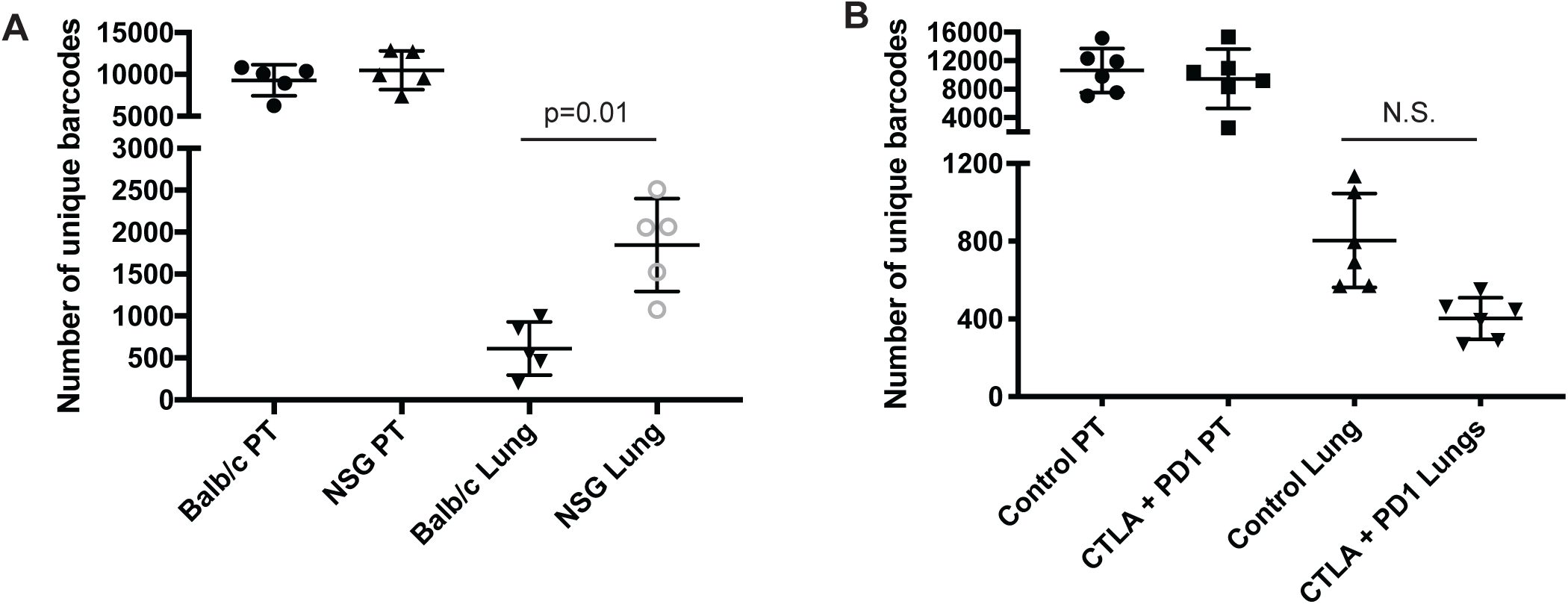
Analysis of changes in barcode proportions in the 4T1 cells with the 300 000 barcode library replicates the 5000 barcode library. A. Number of unique barcodes identified in 4T1 primary tumours and lung metastases grown in NSG mice or Balb/c mice. GLM fit with Tukey’s HSD for multiple comparisons. n= 5 mice per group. B. Number of unique barcodes identified in 4T1 primary tumours and lung metastases grown in Balb/c mice treated with isotype control antibodies or anti-PD1 + anti-CTLA4. A trend to decreasing unique barcode number is seen in anti-PD1 and anti-CTLA4 treated lungs, although this does not reach significance. GLM fit with Tukey’s HSD for multiple comparisons. 5-6 mice/group.

**Sup Figure 9.**
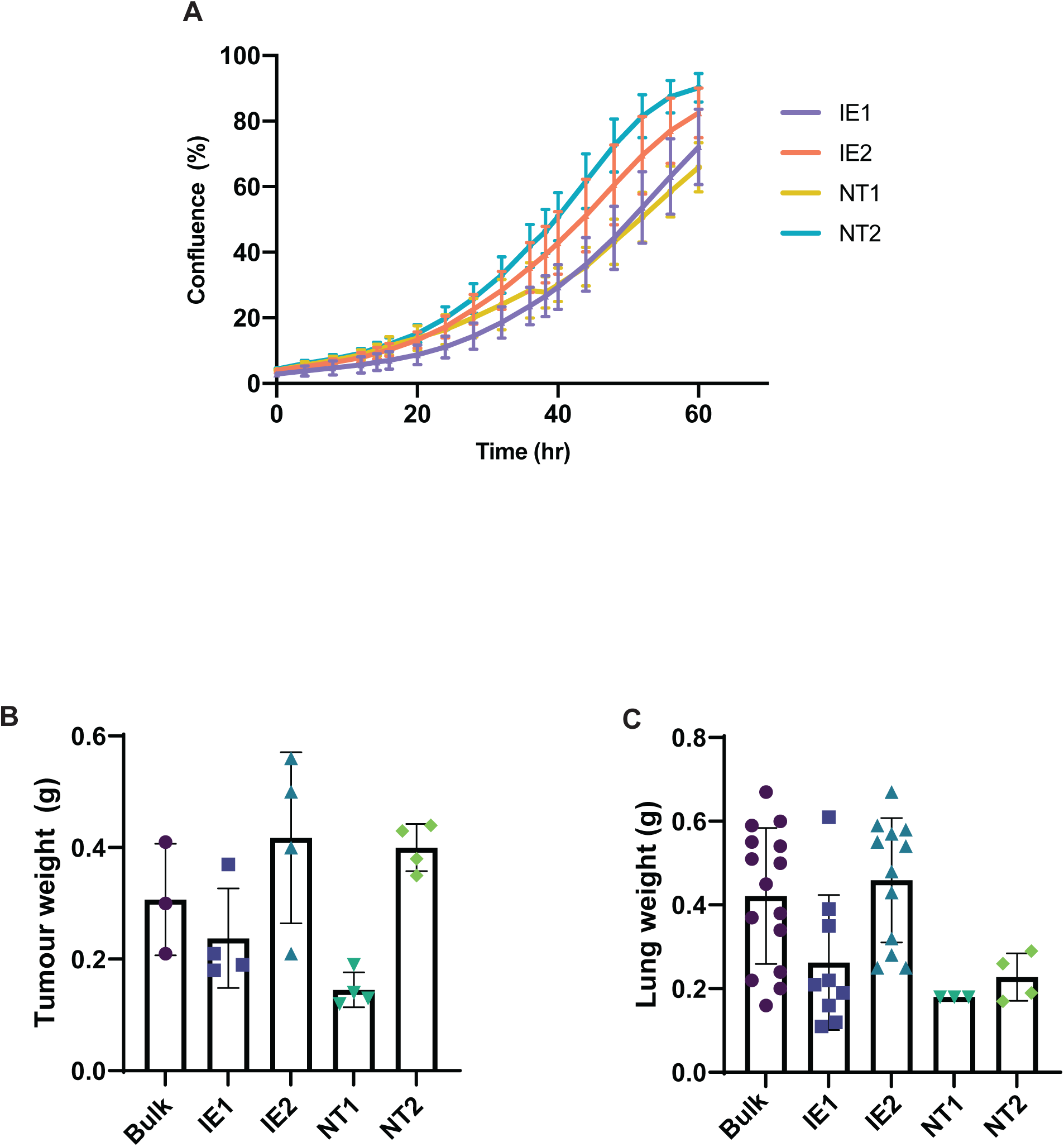
Kinetics of 4T1 clonal cell lines in vitro and in vivo. A: Growth kinetics in vitro as measured by percentage confluence over time (hr). B: Tumour weight at resection generated from clonal cell lines transplanted into BALB/c mice. C: Lung weight at endstage of untreated subclone-tumour bearing BALB/c mice

**Sup Figure 10.**
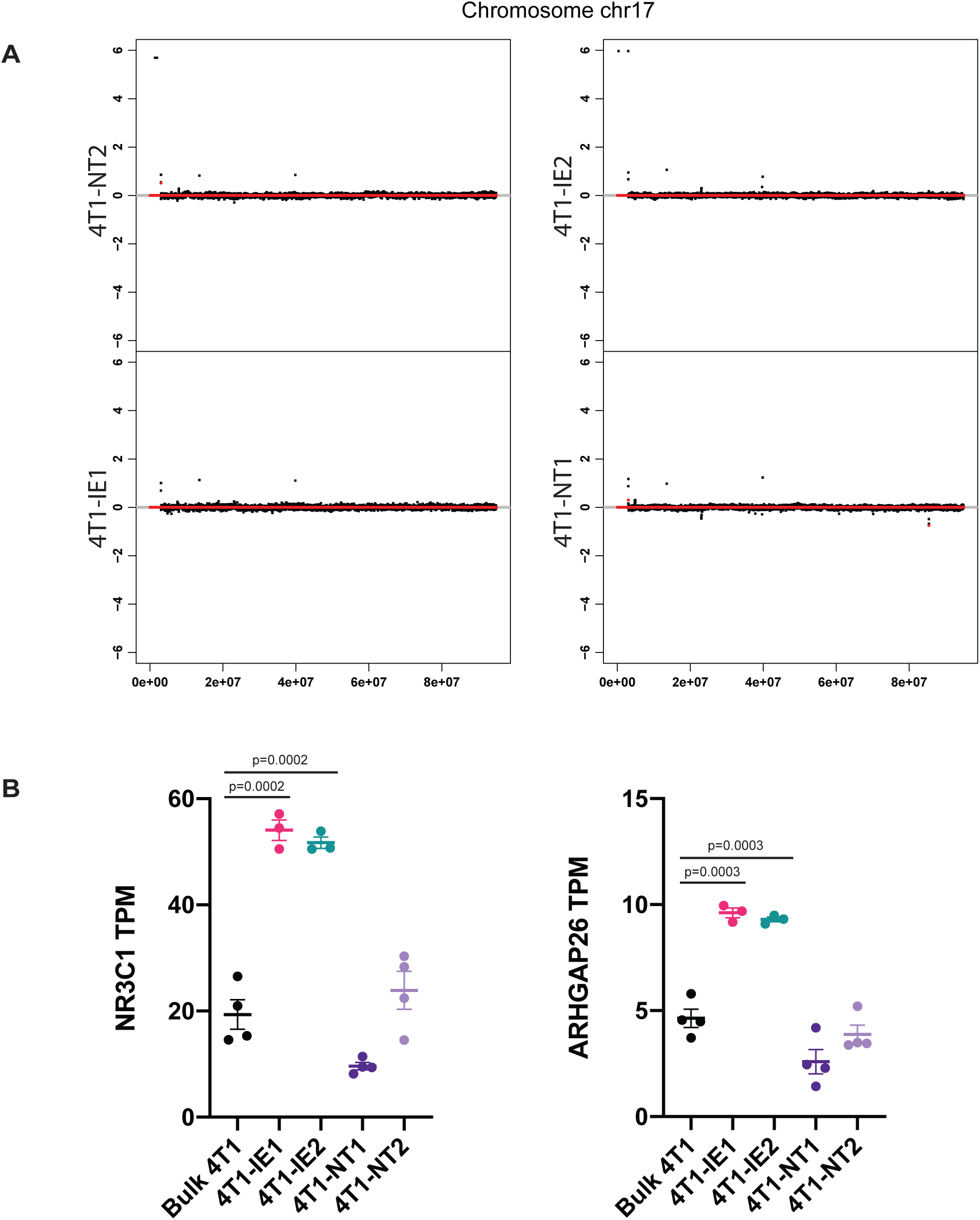
Genomic alterations do not appear to explain phenotypes of the immune evasive clones A. Genome copy number at the MHC-I locus on chromosome 17 of indicated clones. Chromosomal location is indicated on the x-axis and level of copy number alteration is indicated on the y-axis. B. Expression levels of the two genes that have a single copy number increase in IE1 and IE2 in indicated cell populations.

**Sup Figure 11.**
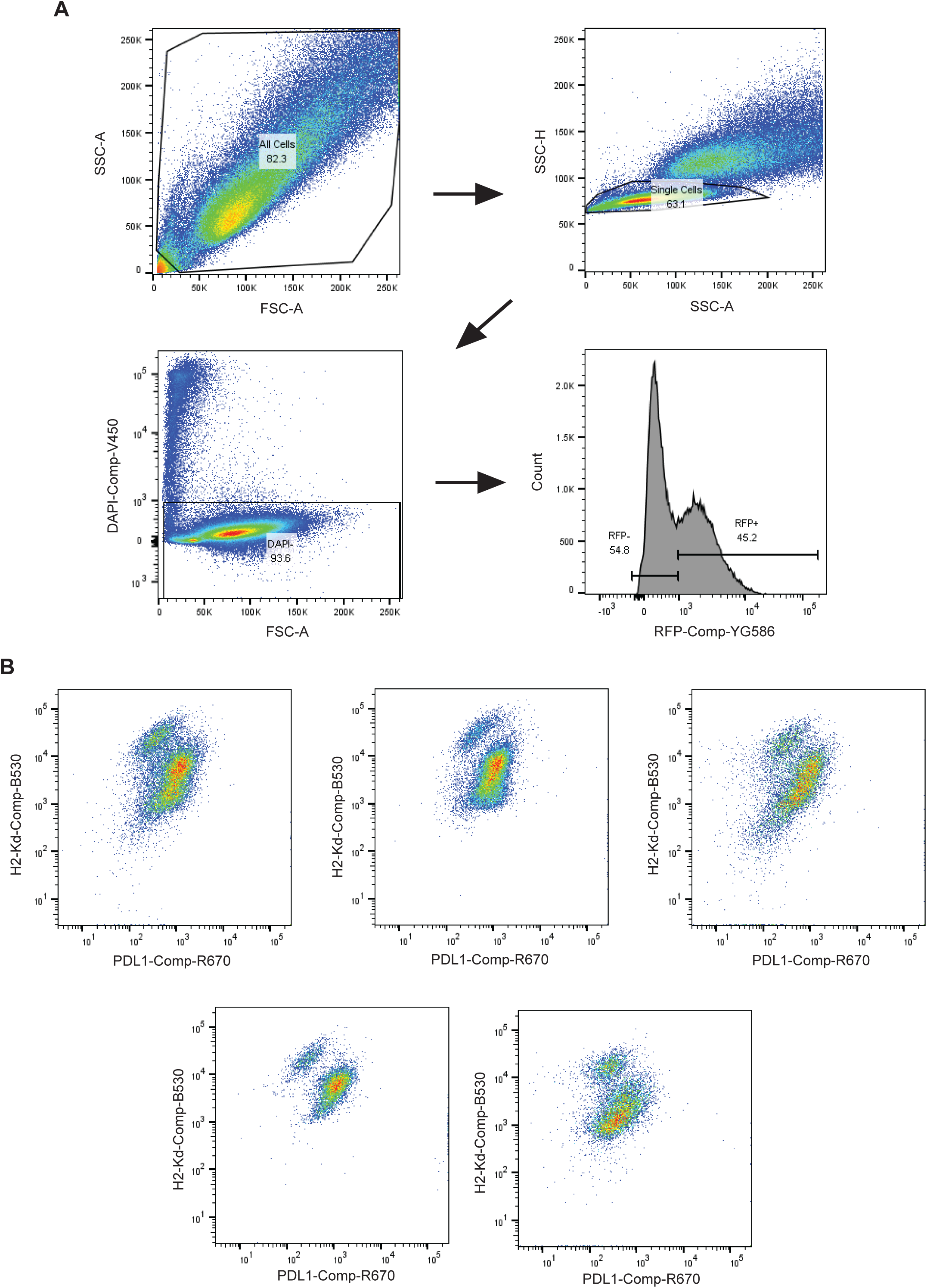
Distinct populations of subclones with varying PD-L1 and MHC I expression exist in immunotherapy treated end-stage lungs. A: Gating strategy. B: PD-L1 and MHC I expression of RFP+ barcoded cancer cells isolated from endstage 4T1 immunotherapy treated lungs. Each plot represents an individual mouse.

**Sup Figure 12.**
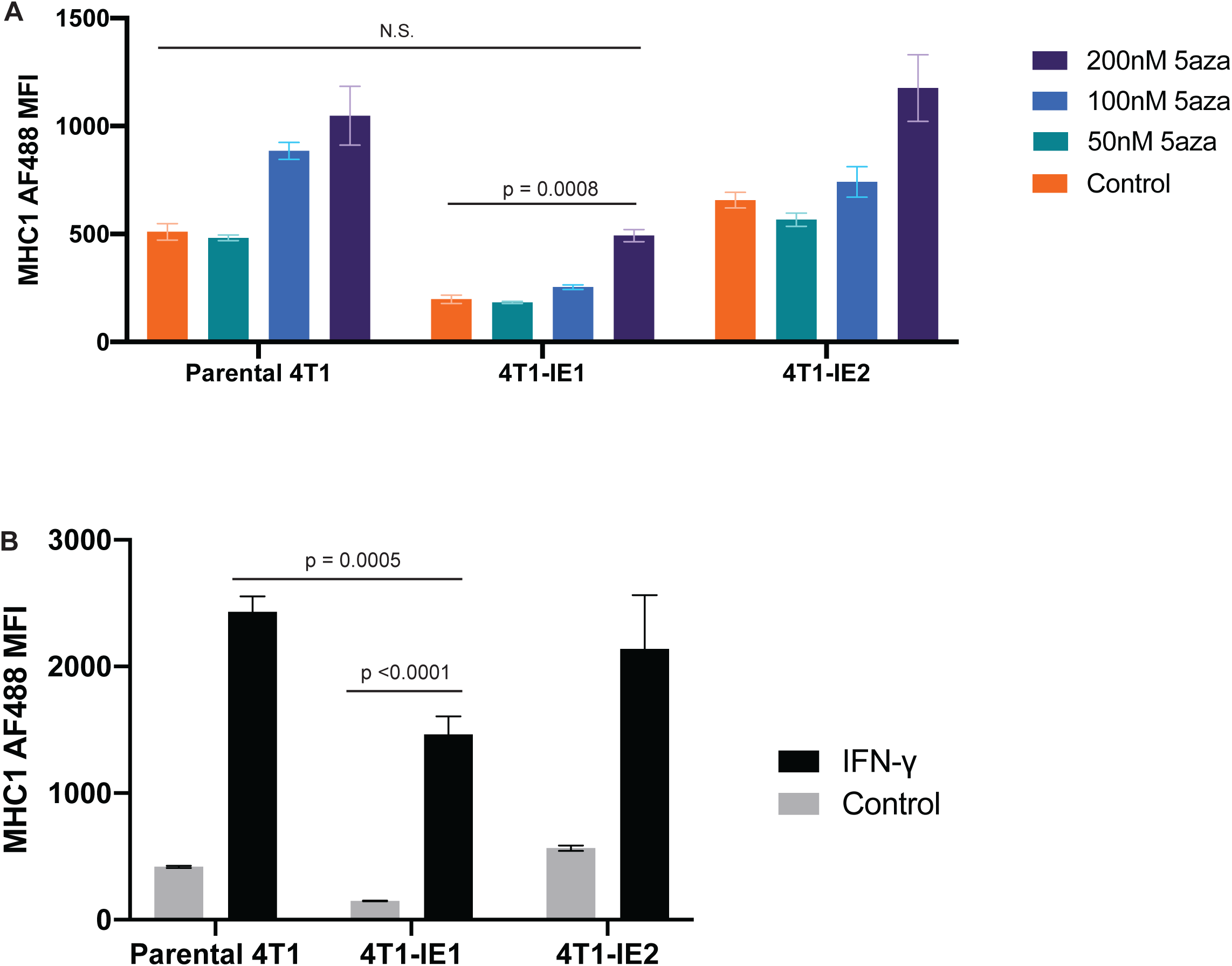
Regulation of MHC expression by 5-aza and IFN-γ in the clonal cell lines. A. MHC-I protein expression measured by flow cytometry in indicated cell lines treated with indicated concentrations of 5-aza. Two-way ANOVA with Tukey’s HSD for multiple corrections. B. MHC-I protein expression in indicated cell lines without or with IFN-γ treatment. One way ANOVA with Tukey’s HSD for multiple corrections.

**Sup Figure 13.**
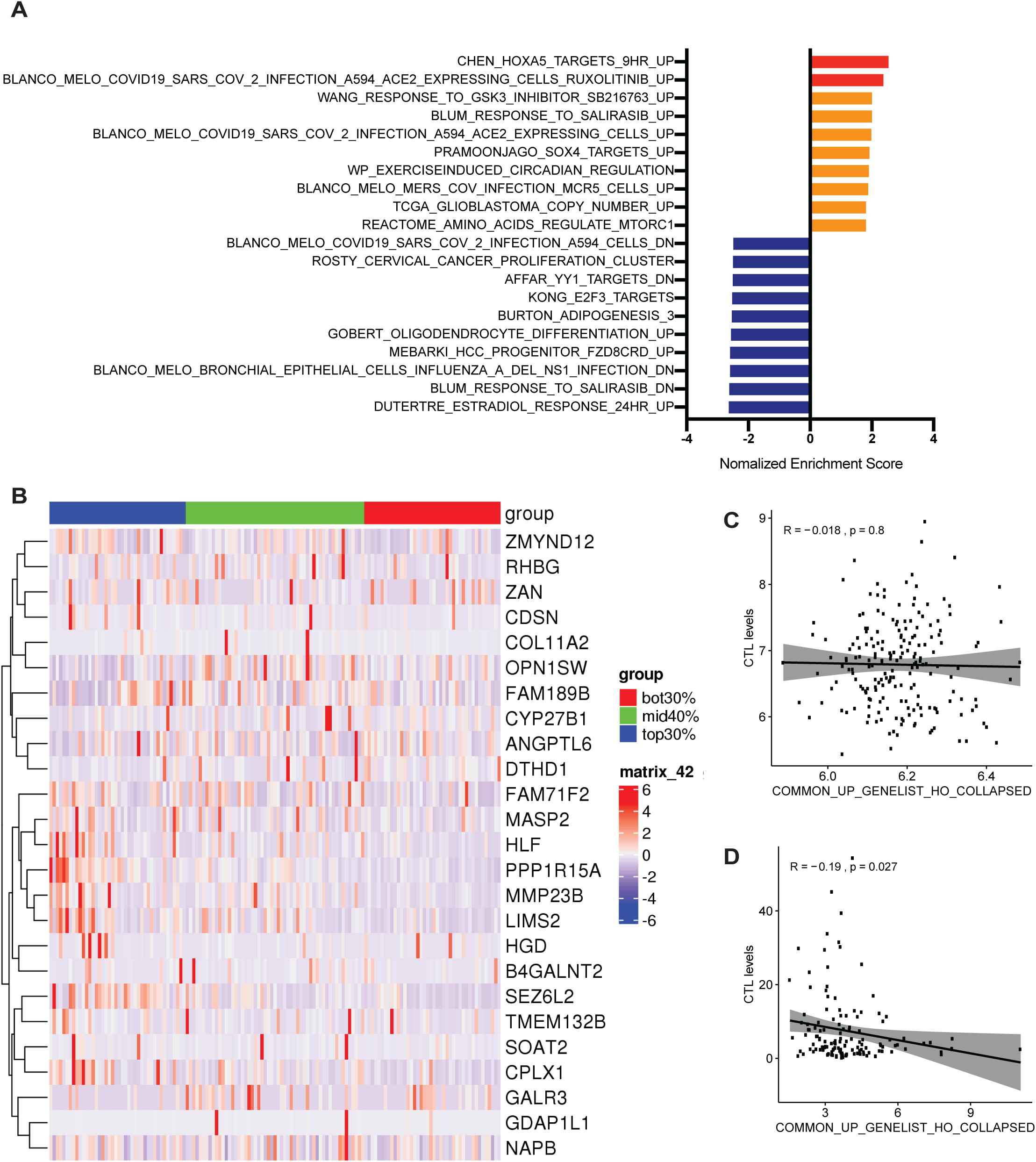
Common gene signature from immunotherapy resistant clones. A. Gene-set enrichment analysis of the overlapping genes between the IE1 and IE2 clones. Red indicates upregulated gene-sets with a significant FDR q-value when multiple testing is taken into account, orange indicates significant upregulated gene-sets with a nominal p value, and blue indicates downregulated gene-sets with a significant FDR q-value. B. Heatmap of unsupervised hierarchical clustering of the immunotherapy resistance signature genes in the TCGA breast cancer dataset, with tumours grouped based on top 30%, middle 40%, and bottom 30% overall gene signature score. C. Correlation plot of immunotherapy resistance signature score with cytotoxic T lymphocyte (CTL) levels in the METABRIC dataset. D. Correlation plot of immunotherapy resistance signature score with CTL levels in the TCGA dataset.

Sup Figure 6: Depletion of CD8+ T cells with anti-CD8, CD4+ T cells with anti-CD4 or NK cells with anti-asialo GM1 does not significantly affect survival of mice bearing 4T1 tumours. A. Kaplan-Meier survival analysis of Balb/c mice transplanted with 4T1 cells, primary tumour was resected on day 15. Depletion of target cell types was initiated 1 day prior to resection, on day 14. 8-15 mice per treatment group. B-D. Flow cytometry to confirm depletion of target cell types. Buffy coat was collected on day 22, 2 days after the final dose of depleting antibodies was given. One way ANOVA.

## Supplementary Table Legends

Sup Table 1: Barcode overlap between two cell pellets from independent barcoding experiments using the 4T1 barcoded cell line. At a barcode cutoff of 5, more than 95% of barcodes detected were the same, and at a cutoff of 10, more than 97% of detected barcodes were identical.

Sup Table 2: DNA barcode insertion sites of IE1 and IE2. By searching the whole genome sequencing (WGS) data and identifying read pairs where only one read mapped to the barcode sequence, the matching mate was blasted against mm10 to identify the barcode plasmid insertion site. Both barcodes were found to reside in introns of cancer-related genes but there was no difference in expression of these genes detected.

Sup Table 3: Copy number variation (CNV) locations found within all subclones. By analysing whole genome sequencing data in R using the cn.mops pacakges, copy number variations could be determined. No major copy number aberrations were detected across the clones, although a single copy number gain was detected in IE1, IE2 and NT2.

Sup Table 4. List of genes differentially expressed in IE1 compared to parental 4T1 bulk. Differentially expressed genes (DEGs) were generated by analyzing bulk RNA sequencing data using R and the EdgeR package. DEGs were filtered for significance based on a FDR <0.05.

Sup Table 5. List of genes differentially expressed in IE2 compared to parental 4T1 bulk. Differentially expressed genes (DEGs) were generated by analyzing bulk RNA sequencing data using R and the EdgeR package. DEGs were filtered for significance based on a FDR <0.05. A greater number of differentially expressed genes were detected in IE2 than IE1.

Sup Table 6. All significantly enriched gene sets found in IE1. Differentially expressed genes generated from comparing IE1 to bulk were preranked by fold change before searching for gene set enrichment using the Molecular Signature Database (MSigDB) across all available collections. Gene sets were filtered for significance based on a FDR<0.05.

Sup Table 7. All significantly enriched gene sets found in IE2. Differentially expressed genes generated from comparing IE2 to bulk were preranked by fold change. Gene set enrichment was carried out using Molecular Signature Database (MSigDB) across all available collections. Gene sets were filtered for significance based on a FDR<0.05

Sup Table 8: List of common differentially expressed genes found in IE1 and IE2. Commonly differentially expressed genes (DEGs) were determined by overlapping significant DEGs in IE1 and IE2 by gene name. An average fold change was calculated across the two samples.

Sup Table 9: Top significantly enriched gene sets found from common differentially expressed genes in IE1 and IE2. The fold change of the common differentially expressed genes in IE1 and IE2 were averaged together to generate an average fold change across both IE1 and IE2. The gene list was then preranked before searching for gene set enrichment using Molecular Signature Database (MSigDB) across the C2 All collection. Gene sets were filtered for significance based on a FDR<0.05. The majority of significant gene sets that were returned were negatively enriched in IE1 and IE2.

